# The dynamic transcriptome of *Plasmodium malariae* in the mosquito

**DOI:** 10.1101/2025.05.05.652270

**Authors:** J. L. Ruiz, B. Díaz-Terenti, T. Lefèvre, R. S. Yerbanga, T. D. Otto, E. Gómez-Díaz

## Abstract

To survive and multiply in different host niches, malaria parasites require sets of proteins that are expressed during the life cycle in a timely manner. *Plasmodium malariae* is a human malaria parasite that is present in most malaria-endemic regions. However, this species remains enigmatic because it is very challenging to culture, its low parasitaemia in human infections, and frequent co-infection with other malarias. We investigate the transcriptome of *P. malariae* during transmission in the vector by analyzing RNA-seq data from midgut and salivary gland parasite stages obtained in two experimental mosquito infections with field *P. malariae* isolates. Our analysis resulted in 3,699 expressed genes, of which 263 are developmentally regulated and 1,338 are *P. malariae* specific, including genes with unknown functions and without orthologs in other *Plasmodium*. We detected unique expression patterns of the ApiAP2 family of transcription factors, many of which appear to function as master regulators during sporogony. We found expressed several members of multigene families, like PIR, PHIST, fam-l, and described new families potentially showing transcriptional heterogeneity. Our findings point to the uniqueness of the *P. malariae* transcriptome, probably related to different transmission traits in the vector, and the lower pathogenicity and virulence in the human.

## INTRODUCTION

Human malaria is a devastating disease and the cause of a huge global health burden, with 263 million cases of disease and 597,000 related deaths in 2023 (1). It is caused by six species of protozoan parasites of the genus *Plasmodium*. *P. falciparum* is the most prevalent and virulent, and, together with *P. vivax*, are responsible for the majority of the morbidity and mortality (1–3). However, there are other species currently unacknowledged, for example, *P. malariae* which is present in most malaria-endemic regions, is found in at least 4-24% of the human malaria cases (4–10). The prevalence in sub-Saharan Africa ranges from 0 to 32% and up to 66% in some countries (10–12). Importantly, the prevalence is thought to have significantly increased over time (13).

The true burden of *P. malariae* and the contribution to global malaria transmission and epidemiology is currently unknown due to the less abundant diagnosis and the paucity in the collection of data for this species (14). What we know is it causes milder clinical manifestations, with lower numbers of merozoites and parasitemia, and moderate recurrent fevers (15,16). Despite this lower virulence and morbidity, *P. malariae* can still cause prolonged and subpatent malaria infections, even for a lifetime (17,18), that may lead to anemia, adverse pregnancy outcomes, chronic renal dysfunction and death (15,19–21). As coinfecting pathogen, *P. malariae* may also modulate the clinical course and transmission patterns of other Plasmodium spp.: it can favor *P. falciparum* infections (22–26), it has been shown to be less susceptible to common and novel antimalarial compounds (27–34), and in consequence it may thrive in regions where *P. falciparum* is declining due to malaria control approaches (5,35–37).

Knowledge about the biology and the life-history characteristics of this neglected pathogen is also very limited. The development in humans differs from the other human malarias in a longer intraerythrocytic cycle (IDC) of approximately 72 hours and the ability to relapse (15). Prepatent period has also been shown to vary, ranging from 16 to 59 days depending on the *P. malariae* strain (15). As for the sporogonic cycle, eight *Anopheles* species have been reported to be competent vectors (15), which is a low number compared to the approximately 70 species known to transmit *P. falciparum* (38,39). Development within the mosquito (i.e. sporogony) is also distinct from *P. falciparum*, which develops for 12-14 days (40,41), while *P. malariae* can take longer, 14 to 21 days (15,21). Similar to other malaria parasite species, the exact duration of the sporogonic cycle or the number and infectivity of the sporozoites have been shown to be highly variable and dependent on several mosquito intrinsic and extrinsic factors (14,40).

These phenotypic differences between *P. malariae* and other malaria parasite species are expected to reflect distinct transcriptomes probably associated to species-specific gene repertoires. The publication of the *P. malariae* reference genome revealed that this species has the largest genome of all human-infecting Plasmodium which is partly due to the presence of large species-specific families or lineage-specific expansions (42–44). Many of these correspond to multigene families located in telomeric and sub-telomeric regions. Indeed, in *P. malariae*, around 40% of the genome is sub-telomeric (44), compared to ∼10% in *P. falciparum* (45). Apart from this, there is also a large number of genes of unknown function lacking functional annotation in *P. malariae*. Alternatively, the divergent transcriptome could be related with differences in how common sets of genes are transcriptionally regulated. This phenomenon has been reported for example when comparing *P. falciparum* and *P. vivax* genes encoding MSP and RBP proteins (46). However, because of the paucity of transcriptomics and epigenomics data in *P. malariae*, no evidence exists for differences in regulation compared to other species.

*P. malariae* contains a specific repertoire of the ApiAP2 family of transcription factors. The role and significance of ApiAP2 TFs for the development and survival of *Plasmodium* is being increasingly recognized, and several ApiAP2 proteins have been described to be master regulators in various life-stage transitions (29,47,48). Of the 29 AP2 proteins that have been annotated in *P. malariae*, only 7 are orthologues (6 also syntenic) to *P. falciparum* AP2 TFs (49–51), while 15 are ortholog to one or more rodent *P. berghei* AP2 encoding genes. The fact that nearly half of the repertoire of AP2 TFs, at least 13 of these proteins, are *P. malariae* species-specific, could be pointing to unique functions. However, the expression patterns and function of AP2 proteins in *P. malariae* are mostly unknown.

The presence of novel and expanded multigene families that are located in subtelomeric regions is indeed a distinctive feature of *P. malariae*. Examples include the *pir* gene family, present in all human malaria parasites except in *P. falciparum* (52,53), the STP1 family, which is expanded in *P. malariae*, and two novel multigene families organized in tandem termed *fam-m* and *fam-l* (formerly *fam-a*) (44,54). Interestingly, the *fam-m* and *fam-l* families do not have orthologs in any other species, but they display structural similarity to a *P. falciparum* merozoite surface protein known to be essential for red blood cell invasion, Rh5 (44,55). Despite most of these specific families are likely involved in host-parasite interactions, virulence and pathogenesis, their transcriptional dynamics and biological functions in *P. malariae* still remain understudied. Further, because of the dearth of genomics and functional studies, it is not known whether additional gene families (multigene or not) display transcriptional heterogeneity in *P. malariae* and thus could be defined in this species as clonally variant (or hypervariable in the context of single-cell transcriptomics studies).

Gene expression regulation during development has been extensively studied in blood-stages of *P. falciparum* (56–63) and *P. knowlesi* (53,64–71). For *P. malariae*, transcriptomics data is scarce: 1) two single-cell RNA-seq datasets for blood stages within the Malaria Cell Atlas with only ∼300 cells, and 2) a bulk RNA-seq study that investigated *P. malariae* transcriptome in the context of an experimental human infection. For the later, data partly corresponded to the gametocyte stage, but other intraerythrocytic stages were present, and due to the very low amount of mRNA, expression was detected just for ∼20% of the genes (72). This paucity of data is partly due to lower parasitaemia levels of *P. malariae* in the blood in human infections, and because for this species *in vitro* culture has not been possible so far. All the above make the mosquito an ideal model system to study transcriptional regulation in *P. malariae*.

Transcriptomic data for the mosquito life-cycle is available for some human- and rodent-infecting *Plasmodium spp.*, including *P. falciparum* and *P. vivax*, and *P. berghei*, respectively. Previous studies have reported just-on-time coordinated waves of gene expression controlling major developmental transitions during sporogony (63,73–79). In relation to the AP2 family of TFs, ApiAP2-encoding genes display a temporal dynamic in gene expression during sporogony similar to the one reported for the blood cycle, but the TFs are distinct and mosquito stage-specific. A few master regulators have been identified controlling expression cascades of the oocyst and sporozoite stages, chiefly is PfAP2-EXP TF with binding sites in the promoters of hundreds of sporozoite-specific genes (76,80,81). As for multigene families, earlier work in *P. falciparum* has shown that the mosquito passage activated the expression of several clonally variant genes with key functions in virulence and pathogenesis in the human but unknown functions in the mosquito host, including *var* genes (82). Whether these patterns are more or less different in *P. malariae*, reflecting differences in virulence and transmissibility, is not known.

In this study, we aim to characterize the transcriptome of *P. malariae* during its development in the mosquito. To this end, we conducted a series of experimental infections of *An. gambiae* mosquitoes using *P. malariae*-infected blood from human donors and performed RNA-seq of mosquito midguts and salivary glands at 16 and 18 days-post-infection (DPI). For the first time, we identified species- and stage-specific transcriptional programs in this pathogen during sporogony, including several genes of unknown function, and unique expression patterns of the ApiAP2 family of regulatory proteins. We also describe the expression of known multigene families implicated in parasite virulence and immune evasion, and genes/families showing evidence of transcriptional heterogeneity in other Plasmodium *spp*.

## MATERIAL AND METHODS

### Mosquito rearing and experimental infections

Three- to 5-day-old female *An. gambiae* s.s. laboratory mosquitoes, sourced from an outbred colony established in 2008 and repeatedly replenished with F1 from wild-caught mosquito females collected in Soumousso, near Bobo-Dioulasso, southwestern Burkina Faso (West Africa), were used for experimental infections. Mosquitoes were maintained under standard insectary conditions (27 ± 2 °C, 80 ± 5% relative humidity, 12:12 LD). The detection of *P. malariae* parasites in the blood of malaria-infected patients was performed by light microscopy. Two independent experimental infections, biological replicates, were carried out by membrane blood feeding in the laboratory as described previously (83,84). Briefly, females were fed through membranes on *P. malariae* gametocyte-infected blood. Thereafter the mosquitoes were given access to a solution of 5% glucose/ 0.05% 4-aminobenzoic acid *ad libitum*. Dissection of infected mosquito midguts and salivary glands were performed in situ at 16 days post-infection (dpi, blood meal) for midguts and salivary glands of infection 1 and 2, and additionally at 18 dpi for salivary glands of infection 2. Mosquito midgut infection was confirmed microscopically. Immediately after dissection, tissues were stored in TRIzol (Invitrogen) and frozen at -80°C until subsequent processing.

### RNA isolation, RNA-seq library preparation and sequencing

We prepared RNA-seq libraries from *P. malariae*-infected midguts and salivary glands obtained in the two independent experimental infections (Table S1). Total RNA was extracted from pools of ∼J20-30 midguts and ∼50-60 salivary glands using the TRIzol manufacturer protocol. RNA concentration was quantified using a Qubit® 2.0 Fluorometer, and RNA integrity was determined with an Agilent 2100 Bioanalyzer. We used the SMRT-Seq® HT Kit, for the synthesis of cDNA from total RNA, and the Illumina’s Nextera XT DNA Library Prep kit from Illumina, for library preparation, following manufacturer’s instructions. Libraries were sequenced using Illumina NovaSeq 6000 for both 2x250 bp paired-end reads. ∼200 million reads per sample were sequenced on average.

### RNA-seq data processing and gene expression analyses

We assessed the sequences quality and rRNA abundance (using GC content as a proxy) using FastQC v0.11.9 (85). To remove rRNA contamination we used SortMeRNA v4.3.4 (86) with default parameters and the database smr_v4.3_sensitive. If required because different number of paired reads after processing, the script repair.sh from the BBMap v38.97 was used (87).

Because of the occurrence of mixed infections in the field, we first performed a taxonomic classification analysis to assess the presence of other species of *Plasmodium* in the sequencing data, leveraging the unique k-mer counting from KrakenUniq v0.6 (88) for pathogen identification with increased accuracy and sensitivity (89). We used default parameters and the pre-built index kuniq_standard_plus_eupath_minus.kdb.20200816 to include in the analyses the databases archaea, bacteria, viral, human, UniVec_Core, and EuPathDB (eukaryotic pathogen genomes with contaminants removed). To extract only the reads assigned to *P. malariae*, we used the script from KrakenTools v1.0.1 (89) with default parameters, except for-t 5858. These reads were the input in a second round of classification with the same software but this time using a custom database with *P. malariae* and various *Anopheles* reference genomes (PlasmoDB-68 and VectorBase-59, respectively)(90–92). This was performed to further assess and discard the presence of sequencing reads from the mosquito host. Beyond the taxonomic classification approach, a mapping step was still required to discard reads corresponding to other organisms. RNA-seq reads were mapped against the *P. malariae* reference genome using HISAT2 v2.2.1 with default parameters except for --very-sensitive (93). Quality control was performed with Qualimap v2.2.2d (94) and SAMtools (v1.12) (95). We extracted paired unique reads with samtools view -q 60 -f 0x2 and removed sequencing duplicates using picard MarkDuplicates v2.24.0 (96) with default parameters except for --REMOVE_SEQUENCING_DUPLICATES true. To ensure proper alignment and removing noise, we also filtered out reads that aligned less than 100 bp of length by parsing the CIGAR string with basic functions. Correlation and PCA plots were performed by the deepTools2 (v3.5.1)(97) functions multiBamSummary, plotCorrelation, and plotPCA.

After the alignment and quality control steps, raw and TPM-normalized counts at the gene level were obtained using CoCo (v.0.2.6) (98). Default parameters were used except for determining strandedness (-s 0) and providing the insert size average for each sample (-i). The *P. malariae* annotation was adapted beforehand using basic functions, to add the biotype to the gene entries and to extend their 5’ ends by 1.5 Kbp so 5’UTRs were accounted for. Genes with < 25 TPM counts when summing all samples were excluded. We categorized normalized counts in high, medium or low groups dividing the signal in three quantile groups using the *Hmisc::cut2* R function (99).

We assessed the potential multiclonality of the infections by SNP calling using BCFtools v1.14 (100). We used bcftools mpileup and bcftools call with default parameters except for -cv - Ov --ploidy 1, we discarded indels, applied a quality threshold of 30 with VCFtools v0.1.16 (101), and filtered took high quality genotypes (DP > 5). We then manually computed alelle frequency: ref_total = DP4[1] + DP4[2]; alt_total = DP4[3] + DP4[4]; AF = alt_total/(ref_total + alt_total).

To assess the presence of mixed parasite developmental stages in the midgut and salivary gland samples, we first integrated our data with that of other Plasmodium species using the corrplot function and Pearson correlation. Alternatively, we performed digital deconvolution using the CIBERSORTx webserver (102,103). Signature matrices were built using *P. falciparum* and *P. berghei* scRNA-seq data from the Malaria Cell Atlas initiative (104,105) and cell fractions in each bulk sample were imputed using 1:1 orthologs between species. Default parameters were used except for batch correction with S-mode using the same scRNA-seq reference matrix, quantile normalization enabled, and 500 permutations performed.

Differential gene expression analyses were conducted using the DESeq2 R package v1.30.0 (106). Correlation and PCA plots by deepTools2 showed higher similarity between infections than between stages (Fig S1A-B). Accordingly, experimental infections were used as replicates and included as a covariate in the formula. The significance threshold was established at p.adj (adjusted P-value) < 0.05. Gene Ontology (GO) enrichment analyses were performed by PlasmoDB tools (default parameters, P-value threshold 0.05). Unless otherwise specified, we used R for basic statistical and data mining analyses (107) and the packages iheatmapr (v0.5.0) (108) and ggplot2 (v3.3.3) (109) for plotting results. The genome browsers used to assess genome-wide transcriptional profiles, motif instances, and gene features was IGV (110). Coverage profiles were converted to bigwig files normalized by bins per million mapped reads (BPM) using deepTools2 (97) bamCoverage function.

### Identification of putative novel multi-copy variable gene families in *P. malariae*

We aimed to identify *P. malariae* genes encoding protein products with similar or identical functions, which could serve as a proxy to expect some degree of heterogeneity in their transcriptional patterns (i.e., HVGs). Using PlasmoDB-68 annotation, we focused on the ’product description’ annotation for all *P. malariae* genes and extracted those with identical values. Subsequently, genes with very generic descriptions, such as ’hypothetical protein’, ’conserved Plasmodium protein, unknown function’, ’conserved protein, unknown function’, and genes belonging to clonally variant multigene families previously described by other authors (44,82) were eliminated. Next, a manual search was performed leveraging PlasmoDB visualization to determine whether genes sharing a description were also located in tandem. Finally, beyond the semantic similarity of their annotated product, we assessed the degree of sequence homology between all the members of the gene families. To this end, we first extracted the nucleotide sequences via PlasmoDB interface and used blastn v2.16.0 (111) with default parameters except for -evalue 0.0001. To establish a reference threshold, we computed the intra-family homology of the families of well-described multi-copy and clonally variant gene families, namely fam-l, fam-m, pir, stp1, clag, rbp, trag. We report ∼80% as the average identity in these families and, together with an identical description of the coding product, an identity above this threshold was set as a requirement to be considered as a multigene family candidate in this study.

As an alternative strategy, we analyzed external single cell RNA-seq data to identify *P. falciparum* and *P. berghei* genes showing evidence of cell-to-cell transcriptional heterogeneity in mosquito stages. These were herein named hypervariable genes (HVG). Our assumption was that the ortholog genes in *P. malariae*, if available, could be HVGs also in *P. malariae* and show a pattern of transcriptional variability during sporogony. To identify HVG candidates for each Plasmodium species and developmental stage independently, we extracted for each gene the range of reported expressions in all cells, except for outliers (following the 1.5*IQR rule) to compensate possible dropout events. We categorized all genes with a non-zero range (i.e., non-equal maximum and minimum non-outlier values, so the genes are expressed with variability in multiple cells) into Very High, High, Medium, Low, and Very Low categories (Hmisc::cut2 wiuth g=5 parameter instead of the g=3 above). We consider the genes categorized as Very High/High to be those showing larger cell-to-cell variability of expression. Next, we used the function FindVariableFeatures in the Seurat R package v5.1.0 (112) with default parameters except for selection.method = “vst”,mean.cutoff = c(0.1, 8), dispersion.cutoff = c(1, Inf). As for the parameter “nfeatures”, which is required and determines the arbitrary number of variable features to be obtained based on a standardized variance, we used the number of genes in the Very High/High categories above. We considered high-confidence HVG candidates the genes that are both in the Very High/High groups and in the result of the FindVariableFeatures,and investigated the transcriptional profiles of their ortholog genes in *P. malariae*, if any.

### Genes without annotation in *P. malariae*

Genes with the product description ‘hypothetical protein’, ‘conserved Plasmodium protein, unknown function’, and ‘conserved protein, unknown function’ were considered to be without annotation. To fully unveil potential functions, using PlasmoDB-68 annotation we extracted for these their annotated GO terms, if available. Otherwise, we looked for ortholog genes in *P. falciparum*, *P. berghei*, and *P. vivax*, and interrogated whether there were annotated GO terms in the other species. Ultimately, for the genes without known evidence of function but expressed in our dataset, we applied the PANNZER (Protein ANNotation with Z-scoRE) webserver to obtain automated predictions of protein functions and GO annotations (113,114). The annotations were performed using default parameters, i.e. considering a cut-off value of PPV (Positive Predictive Value) ≥ 0.3, and using the same parameters applied to tripanosomatids in a recent study (115).

### Motif enrichment analysis

We performed *de novo* motif analysis using HOMER v4.11 (116) to the promoters (1 Kbp upstream) of different sets of genes of interest, such as those highly expressed (at least one infection have high expression), to identify the binding sites for *P. malariae* ApiAP2 transcription factors that are ortholog to *P. falciparum* AP2 proteins using the previously published database of validated *P. falciparum* ApiAP2 motifs (117). We used the *findMotifsGenome.pl* module with the parameters “-len 8,10,12 -size given -mset all -dumpFasta” and as reference with the parameter “-mcheck” the *P. falciparum* AP2 motif library file We also conducted motif scan and located the occurrence of motifs using FIMO (118) and the *annotatePeaks.pl* module from HOMER.

### External data

We reanalyzed the RNA-seq bulk dataset available for *P. malariae* blood stages (*i.e.*, mixed stages but correlated with gametocytes) (133) following the same methods applied for mosquito samples in this study.

We downloaded the available scRNA-seq data, corresponding to 13 cells, for *P. malariae* blood stages in the Malaria Cell Atlas (105). We manually computed TPM counts using the functions calculateTPM from the scuttle R package v1.6.3 (119) and reported mean values.

We downloaded from PlasmoDB already published TPM-normalized counts for P. falciparum (75,76,120) and *P. berghei* (121–123) mosquito stages. A dataset of *P. vivax*-infected *Anopheles stephensi* tissues is available (73), but original authors only addressed the mosquito responses and did not analyzed parasite sequences. To obtain TPM-normalized counts for *P. vivax* in mosquito midguts and salivary glands, we reanalyzed this data following the same methods applied for *P. malariae* samples in this study, except for using the *P. vivax Sal-1* reference genome (124,125).

We downloaded scRNA-seq reference data used in the analyses above from the Malaria Cell Atlas, corresponding to *P. falciparum* and *P. berghei* (104,126). In *P. falciparum*, we used cells labeled as “ookinete” (112 cells), “sporozoite-oocyst” (158 cells), or “sporozoite-salivary glands” (257 cells), while in *P. berghei* we used cells labels as “ookinete” (141 cells), “oocyst” (252 cells), or “sporozoite-salivary glands” (153 cells). In all cases above, ortholog genes between *P. malariae* and other Plasmodium species were determined using PlasmoDB and OrthoMCL (127) and only syntenic genes were used except when noted otherwise.

## RESULTS

### The mosquito as an experimental model for studying *P. malariae* infections

RNA-seq libraries were produced from adult female *Anopheles gambiae* s.s mosquitoes fed with *P. malariae*-infected blood obtained from human volunteers from a malaria endemic area in Burkina Faso, Africa (see Methods). Two independent experimental infections were performed; two samples are available for midguts and three for salivary glands (Fig 1A; Table S1). The replicability analysis of the RNA-seq data revealed higher similarity and clustering by tissue rather than by experimental infection, which supports the use of the infections as biological replicates in our transcriptomics analyses (Fig S1A-S1B).

**FIG 1.**
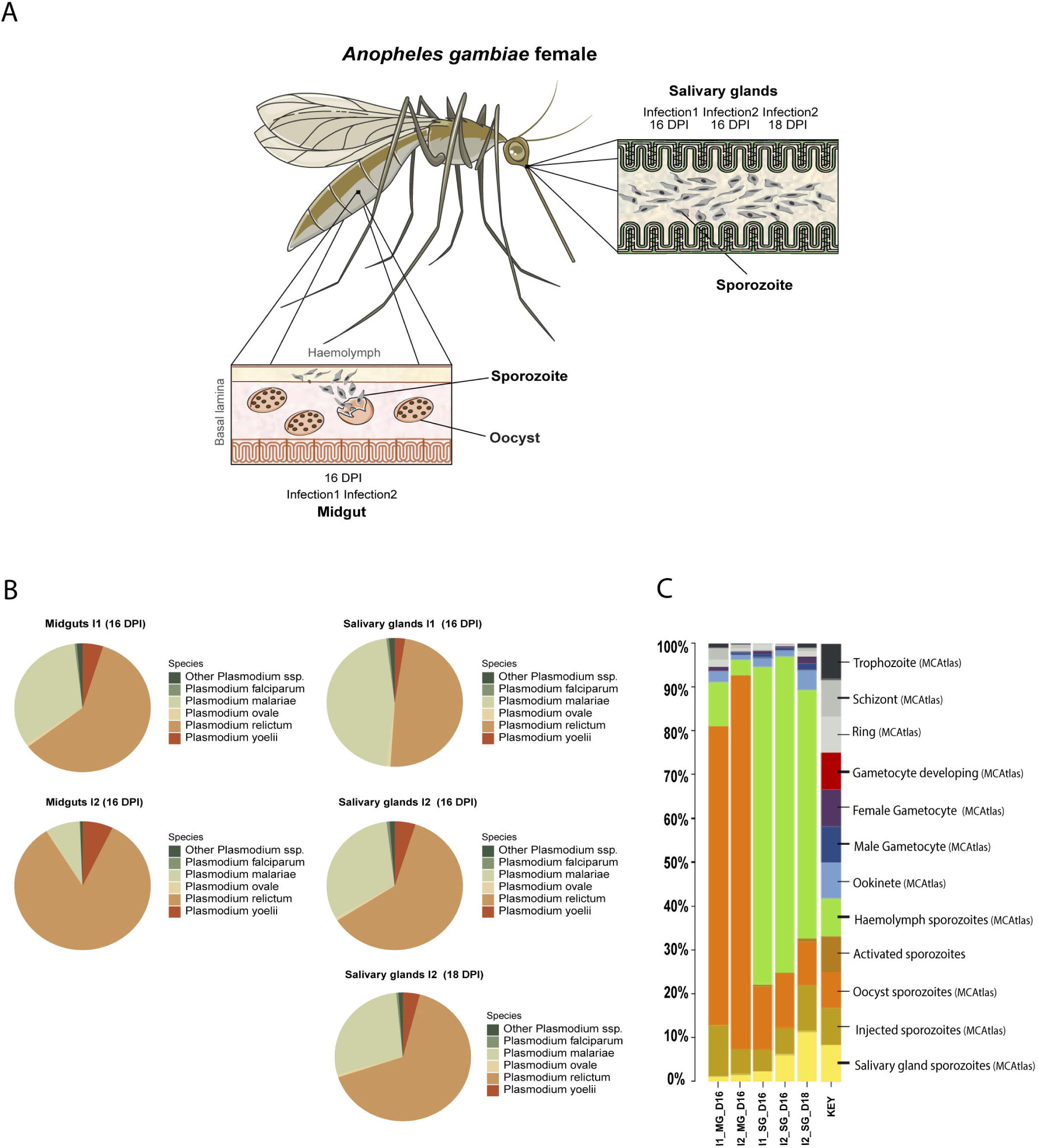
The mosquito as an experimental model to study *Plasmodium malariae* infection. **(A)** Schematic representation of *Plasmodium malariae* development in its natural vector *Anopheles gambiae*. After ingestion of *P. malariae* infected blood, gametocytes differentiate into gametes in the midgut, fertilise to form a zygote that in 24h differentiate into an ookinete that traverses the mosquito midgut epithelium where oocyst formation occurs. Between 14-21 days post infection, the precise timing is uncertain in *P. malariae*, mature oocysts release sporozoites into the haemocoel (haemolymph), from where they migrate to the salivary glands. The sporozoites invade the salivary glands, becoming available for transmission to the vertebrate host during a subsequent mosquito bite. The samples analysed in this study correspond to the different tissues colonized by the parasite during its development in the vector, that is, mosquito midgut and salivary glands containing the oocyst and sporozoite stages, respectively. **(B)** Pie charts showing the proportion of different Plasmodium species detected in the 5 midguts and salivary gland samples from two independent experimental infections (I1; MG and SG -16 dpi, I2: MG 16 dpi and SG 16-18 dpi). Six Plasmodium species were detected: *P. falciparum, P. malariae, P. ovale, P. relictum, P. yoelii,* as well as other species grouped under ‘Other Plasmodium ssp.’. *Plasmodium malariae* is the most abundant of all human infecting species (Table S3). **(C)** Barplot showing the deconvolution of the *P. malariae* bulk RNA-seq data for midgut and salivary gland samples using previously published *P. falciparum* single-cell data (104). Oocyst and sporozoite stages account for the larger proportions.

To use the mosquito as an experimental model, a series of pre-analytical steps and considerations were made. First, we had to assume some uncertainty in relation to the precise developmental stage of *P. malariae* parasites isolated from the midgut of infected mosquitoes. That is, human malaria parasites typically display variable development period in the mosquito midgut (between 7-14 days). It is after 14 to 21 days that the oocyst eventually bursts, releasing sporozoites into the hemolymph that travel to the salivary glands (15). Therefore, achieving precise isolation of the different developmental stages in the infected midguts is challenging (102,128). Further, the development of *P. malariae* in the mosquito remains particularly less characterized compared to other species, so caution is warranted. Due to this uncertainty and the slower extrinsic incubation period (EIP) of this species, in this study infected midguts were dissected at 16-days post-infection (dpi) whereas salivary glands were dissected at 16- and 18-dpi. For these two time points at least one of the stages is expected to be present in either tissue: midgut 16 dpi = oocysts + immature sporozoites, and 16 - 18 dpi = salivary gland mature sporozoites.

At this regards, one potential methodological shortcoming is the possibility that infected midguts contain a mixed population of mature oocysts and immature sporozoites. We compared our *P. malariae* RNA-seq data with time-course data sets from *P. falciparum*, *P. vivax* and *P. berghei*, and looked for significant correlation (Pearson’s, P<0.001). For the midgut’s samples, in *P. falciparum* we reported highest correlation coefficients against immature sporozoites in midguts (∼50%), followed by ookinetes (∼25%) and oocysts (∼15%), whereas in *P. vivax* the peaks of correlation corresponded to early oocysts (∼37%), and in *P. berghei* to ookinetes (∼30%) (Fig S1C). The peaks of correlation for the salivary gland samples coincided with mature sporozoite datasets in all cases (∼75% correlation) (Fig S1C). To further validate the assumption that *P. malariae* reads in midguts mostly captured immature sporozoites in oocysts, we leveraged the *P. falciparum* and *P. berghei* single-cell RNA-seq data (available at the project Malaria Cell Atlas (104,105) to perform digital deconvolution of our *P. malariae* RNA-seq bulk data of mosquito stages (102). We found that immature sporozoites were predicted to make up the largest proportion of cells in the midgut samples (mean ∼77% of “sporozoite (oocyst)” in *P. falciparum* and ∼42% of “sporozoite (salivary gland)” in *P. berghei*; Fig 1C; S1D; Table S4). Accordingly, the majority of cells in samples from salivary glands were also classified as sporozoites (Fig 1C; S1D; Table S4). All this considered, we assumed that our salivary glands samples, either at day 16 or 18 dpi, captured the transcriptome of *P. malariae* sporozoites, while midguts appeared to contain a mixed composition of oocysts and sporozoite stages (oocyst/sporozoites hereinafter).

Second, to explore the possibility that our samples contain mixed parasite infections, we looked for reads mapping all reference Plasmodium genomes. The taxonomic classification analysis (89) show that a large fraction of reads matched *P. relictum,* and secondarily *P. malariae* with a moderate proportion: 32.3% and 8% of plasmodial reads in midguts of I1 and for I2 respectively. In salivary glands, *P. malariae* reads represented 45.4%, 30.9% and 27.6% for I1, I2-16 and I2-18, respectively (Fig 1B; Table S2). Only reads classified as *P. malariae* were kept for further analysis (Table S2). The alignment rates of the cleaned *P. malariae* reads were high, of ∼80% and ∼99% for midguts and salivary glands samples, respectively (Table S3).

A final consideration of this work because of dealing with natural infections is multiclonality (129). To rule out any confounding pattern due to multiple clones being present, we inferred the frequency of homozygous/heterozygous SNPs in our RNA-seq data (see Methods section). Results indicate that despite different SNPs numbers detected, the vast majority of them were homozygous (>90% of SNPs with allele frequency parameter, AF=1), suggesting that the infections in this study were monoclonal (Fig S2A).

### Uniqueness and dynamics of the transcriptome of *P. malariae* during sporogony

The analysis of *P. malariae* RNA-seq libraries resulted in the detection of 3,699 genes that we considered to be expressed (genes with > 25 TPM, see Methods). These represent ∼50% of the reference transcriptome (6,715 genes) (Table S5). A total of 2,586 out of the 3,699 expressed genes corresponded to protein coding genes that we categorized as highly expressed (see Methods) in at least one of the stages in any infection (Table S5). Amongst the top expressed in *P. malariae* mosquito-stages there are genes known to be expressed in other malaria parasites in the same stages of development, such as CSP or SPECT1 (Fig S4A and 2A respectively; Table S5). Other highly expressed genes include gene families specific to *P. malariae*, such as MSP3 or the gamete antigens 27/25 (Table S5).

When comparing *P. malariae* blood and mosquito stages, most genes appear to be unique to a particular stage of development as is shown in Fig 2B where most of the top100 expressed genes in mosquito stages did not appear as expressed in the blood stages. Between midgut oocyst/sporozoites and salivary gland sporozoites, we identify 2,043 developmentally regulated genes with a log2 Fold Change > 2 and with a range up to 10.6 (Fig 2C, Table S5). However, for most of them the change did not reach statistical significance in the differential analyses. That is, only 263 could be considered as significant differentially expressed genes (DEGs): 145 more expressed in midgut samples (oocysts/sporozoites) and 118 in salivary gland sporozoites (Fig 2D; Table S5).

**FIG 2.**
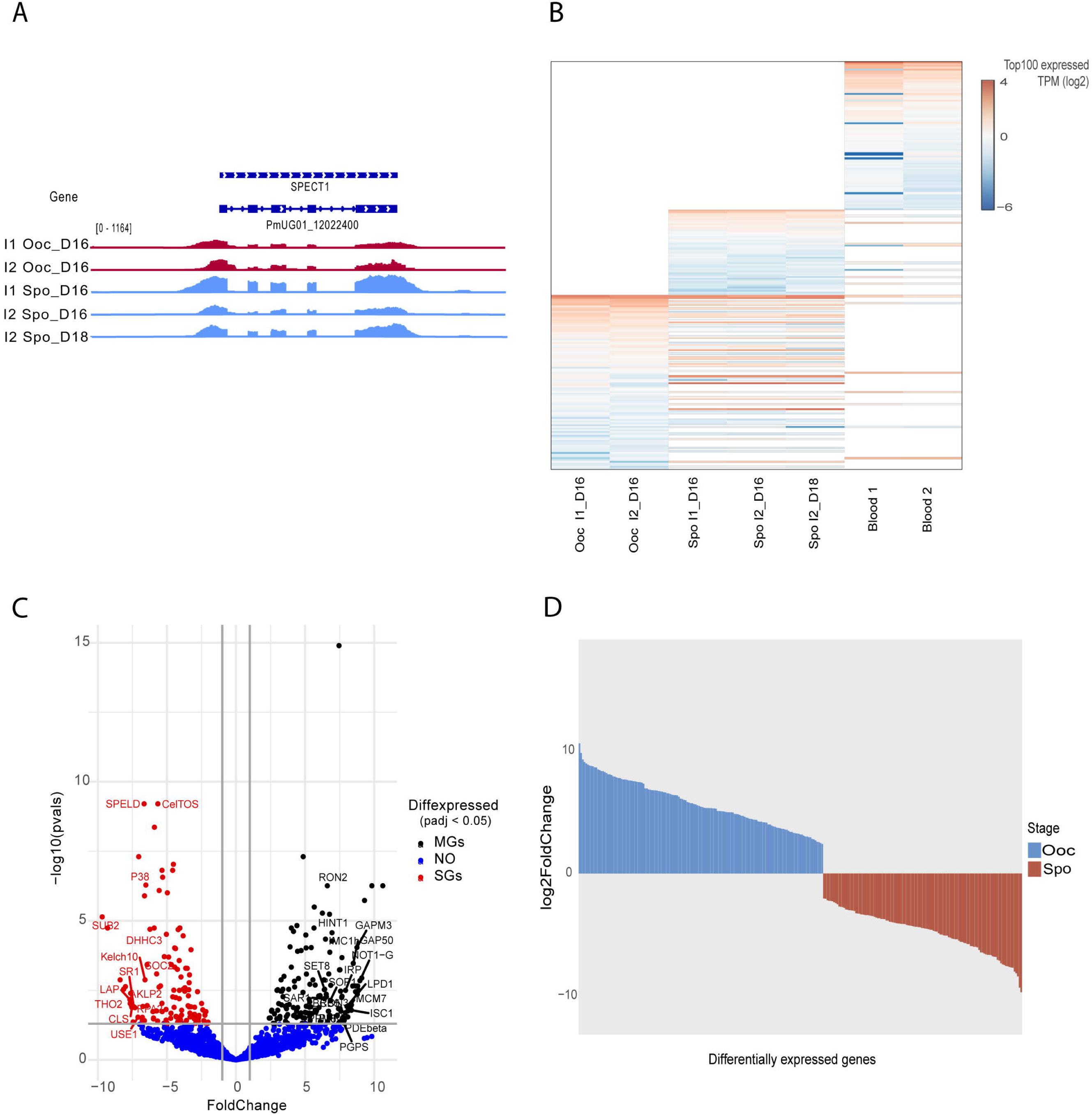
Stage-specific transcriptional profiles of *P. malariae*during sporogony. **(A)** Gene expression profiles in the region containing the gene encoding the SPECT1 gene (PmUG01_12022400). The tracks shown are for midgut oocysts/sporozoites (red) and salivary glands sporozoites (blue). All tracks are shown at equal scale. **(B)** Heatmap for the 100 most highly expressed genes (TPM, log2) in different stages: oocysts/sporozoites (I1_ooc-spo_D16, I2_ooc-spo_D16), sporozoites (I1_spo_D16, I2_spo_D16, I2_spo_D18), and blood (Blood_1, Blood_2). For stages where no expression of certain genes is detected, their expression is blank. **(C)** Volcano plot displaying the magnitude of the change and the statistical significance for differentially expressed genes (DEGs) between *P. malariae* developmental stages. The x axis is the fold change difference between stages, and the y axis shows the level of significance. Significant DEGs are marked in black and red for midgut and salivary glands samples, respectively, the blue ones are not significant. A positive fold change (x axis) indicates higher expression in midgut oocysts/sporozoites, while a negative value indicates the opposite trend (higher gene expression in salivary glands sporozoites). **(D)** Histogram showing the number of differentially expressed genes (DEGs) in midgut oocysts/sporozoites (MGs) and salivary glands sporozoites (SGs).

Our results partially overlap with the dynamic transcriptome during sporogony described in other *Plasmodium* spp. (130–132). For example, DEGs overexpressed in midgut oocysts/sporozoites include inner membrane complex proteins (IMC1a, 1c, 1e, IMC20), the rhophtry protein (RON12), glideosome proteins (GAPM3, GAP50, GAPM2, GAPM1), RALP, TREP, secreted ookinete protein 1 (PSOP1), apical sushi protein ASP or histones (H2A, H2B, H4) (Table S5). Amongst DEGs overexpressed in salivary gland sporozoites, we found the apical membrane antigen AMA1 (Fig S4B), TRAMP (thrombospondin-related apical membrane protein), SPATR (secreted protein with altered thrombospondin repeat domain), SLARP (sporozoite and liver stage asparagine-rich protein), TLP (trap-like protein), 6 cysteine proteins (P38, P52, P36, B9, PmUG01_03014600), PmUG01_07044000 (profilin. motility protein), PUF2 (mRNA binding protein, inhibitor and controls sporozoite conversion to liver stages), PLP1, PLP2 (perforin for host traversal), serpentine receptor SR1, GAMA, the thrombospondin-related anonymous protein (TRAP), the cell traversal protein for ookinetes and sporozoites (CelTOS) or the gamete egress and sporozoite traversal (GEST), sporozoite protein essential for cell trasversal (SPELD) or (Table S5). Similarly, the Gene Ontology overrepresentation analyses on midgut oocyst/sporozoite and salivary gland sporozoite specific genes, showed enriched terms corresponding to functional activities that conform with the expectations for each stage of development (Table S6). These include biosynthetic processes or metabolism in oocysts, and gliding, motility, response to stress or multi-organism symbiotic processes in sporozoites (Fig S2B-C ; Table S6).

Regarding *P. malariae*-specific gene repertoires, a total of 3,347 *P. malariae* genes lack orthologues in the two human malaria species *P. falciparum* and *P. vivax*, or the rodent *P. berghei* species, out of which 1,338 genes appeared expressed in our study. In other words, the mosquito *P. malariae* transcriptome represents ∼40% of the total genes without orthologues and ∼20% of the 6,715 genes in the whole genome. Out of the 1,338 *P. malariae* species-specific genes expressed in the mosquito, 111 are developmentally regulated: 62 of them more expressed in oocyst/sporozoites and 49 in sporozoites, corresponding to ∼3% of the 3,347 genes without orthologues and ∼40% of the total repertoire of DEGs within the mosquito (Table S5). Highly expressed genes within this category include sporozoite specific transcriptional regulators such as ApiAP2 (PmAP2-HS, PmUG01_05037900), TLP, CLAMP and TRAP proteins, which seem to have an important role in *P. falciparum* during the invasion of mosquito salivary glands (133,134). By performing a functional enrichment analysis, the most enriched biological processes (p-value < 0.05) identified include: ‘biological process involved in interaction with host’, ‘biological process involved in symbiotic interaction’, ‘biological process involved in interspecies interaction between organisms’ and ‘movement in host environment’ (Table S7).

Next, we looked at genes without annotation (no gene description) in the *P. malariae* reference genome (see Methods) which represent the ∼20% of the entire transcriptome (1,320 genes) (Table S8). Of these, we report in our dataset the expression of 955 genes (∼70%), and 82 out of the 955 are differentially expressed between midgut oocysts/sporozoites and salivary gland sporozoites in this study (Table S8). To further investigate their possible function, we first considered genes without annotated gene description in *P. malariae* but with GO terms (286 out of the 955 genes), The results of the GO enrichment analysis showed significant terms related to DNA damage and integrity (biological processes) and parasite membrane (cellular components) (Table S9). For the remaining 669 *P. malariae* genes without an associated GO term, we investigated whether they have annotated GO terms in other *Plasmodium* species. To this end, we looked at their ortholog and syntenic genes in *P. falciparum*, *P. vivax* and *P. berghei*. We found 474 orthologs for 343 *P. malariae* genes, and of these 86 orthologs corresponding to 76 *P. malariae* genes have associated GO terms in other species. The functional enrichment analysis revealed GO terms related to the parasite membrane (cellular components, while no significant term for biological processes nor molecular functions) (Table S8-S9).

Finally, on the entire set of 955 expressed genes without annotation we applied a method named PANNZER2 (115) to infer new functional annotations. This analysis resulted in the annotation of 409 genes, of those 136 corresponding to genes that originally did not have any GO term associated. To evaluate the reliability of the new annotations we took the set of genes that had computed and curated GO terms and compared them with those inferred by PANNZER. We obtained 282 matches for 288 genes with both annotations, which corresponds to a 98% of coincidence (Table S8). The new GO terms inferred are in Table S9. The enrichment analysis performed on the expressed genes of unknown function and that have no associated GO terms in *P. malariae* or other *Plasmodium spp*. reveal functions related to proteolysis, symbiont entry to host, adhesion of symbiont host cell (Fig S3A Table S9). The analysis in the set of *P. malariae* genes without annotation but with orthologs, reveals enrichment in GO terms related to symbiont entry to host, adhesion of symbiont host cell and protein processing (Fig S3B-Table S9).

Altogether, our results suggest that the transcriptome of *P. malariae* during sporogony conforms to the just-one-time model described for other Plasmodium spp, however, with a remarkable singularity. That is, the gene repertoire expressed in *P. malariae* mosquito stages is highly species-specific, including hundreds of genes of unknown function, which could play relevant roles in the interaction and adaptation of the parasite to the mosquito and human hosts.

### Master regulators of the *P. malariae* transcriptome during sporogony

The transcription factors from the ApiAP2 family have been shown to act as master regulators of developmental transitions in *Plasmodium spp*., but evidence about their function and essentiality is available only in few reference species, including *P. falciparum*, *P. vivax* and *P. berghei* (49,50,60). Here, we used the mosquito experimental model to find out more about the expression dynamics and target genes of these transcriptional regulatory proteins in *P. malariae*. Out of the full repertoire of 29 AP2 TFs annotated in *P. malariae*, only 7 genes are ortholog (6 also syntenic) to the corresponding *P. falciparum* AP2 TFs, while 15 are ortholog (12 also syntenic) to one or more *P. berghei* AP2 encoding genes, pointing to *P. malariae*-specific functions for at least 13 of these proteins (i.e. without orthologous).

We find that the majority of AP2 TFs (23 out of 29) are expressed during sporogony, although only a few AP2 TFs show noticeable peaks of expression that are consistent across infections (Table S5). In the midgut oocysts/sporozoites, the top expressed AP2 TFs are in this order: AP2-O4 (PmUG01_12014000), AP2-HC (PmUG01_12060900), AP2-EXP (PmUG01_12050500), AP2-O (PmUG01_09052200), AP2-G3 (PmUG01_14033900), AP2TEL (PmUG01_11040700), AP2-L (PmUG01_02024900), PmUG01_09024900, PmUG01_09017400, PmUG01_08023500. In salivary gland sporozoites, the most expressed are in this order: PmUG01_05037900, AP2-EXP (PmUG01_12050500), PmUG01_09017400, AP2-I (PmUG01_08023500), AP2-L (PmUG01_02024900), AP2-O (PmUG01_09052200), AP2-O5 (PmUG01_12067500), AP2-O4, (PmUG01_12014000), PmUG01_09024900 (Fig 3A, Table S5). Remarkably, *P. malariae* AP2-EXP (PmUG01_12050500) is an example of highly expressed AP2 in both midgut oocysts/sporozoites and salivary gland sporozoites (Table S5). The ortholog *P. falciparum* AP2-EXP, was reported to be highly expressed in mosquito stages and potentially involved in transcriptional regulation during sporogony (76). Similarly, PmUG01_09024900, whose ortholog in *P. falciparum* (PF3D7_1115500) is currently of unknown function, appears to be more expressed in mosquito-stages according to published data (75,76).

**FIG 3.**
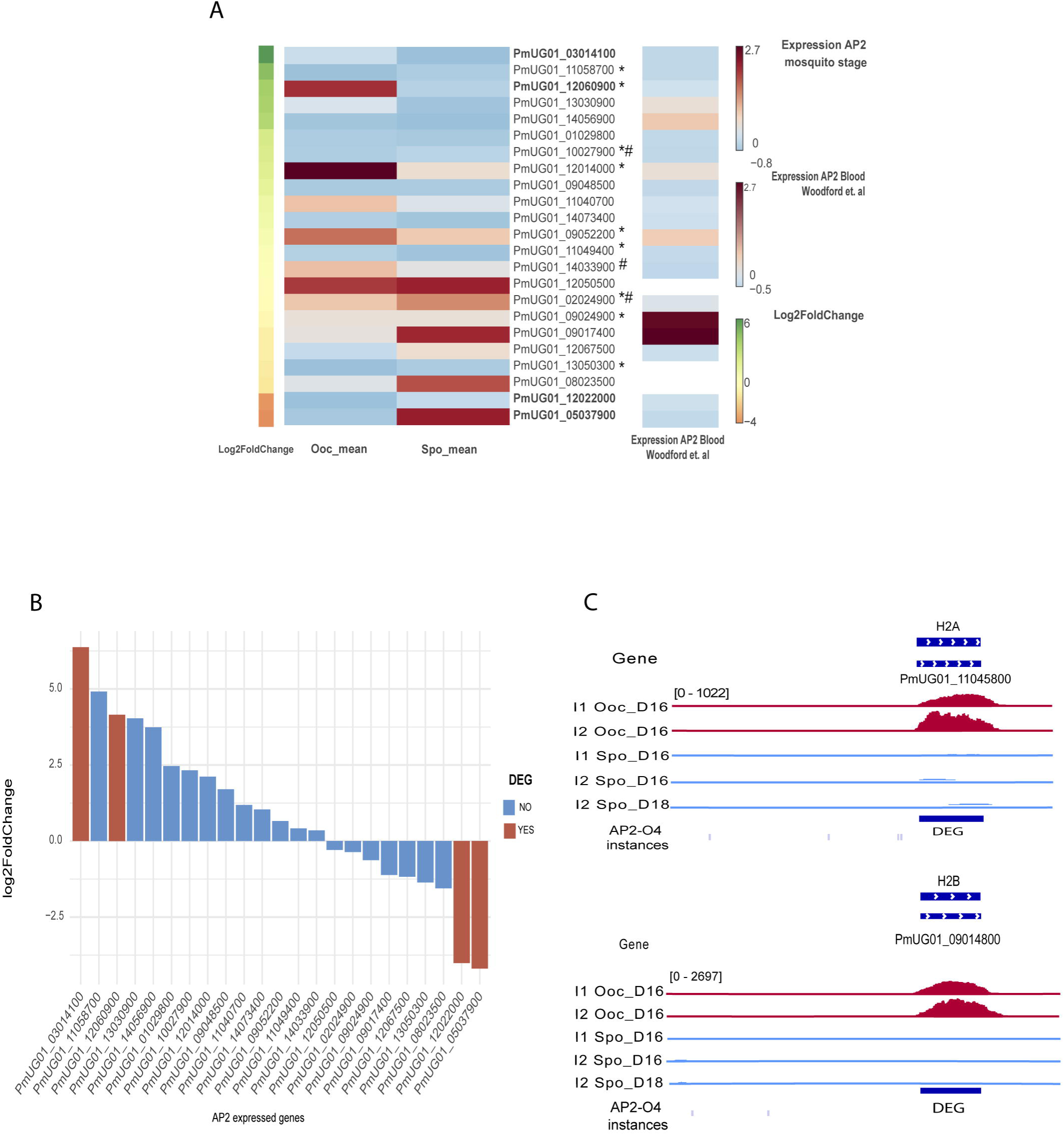
Expression profiles of the AP2 family of TFs in *P. malariae* **(A)** Heatmap of gene expression levels (log2 TPM) for the 29 AP2 annotated proteins in blood and mosquito stages. Differentially expressed genes (DEGs) are highlighted in bold. AP2 orthologues (and syntenic) to *P. falciparum* are shown with (#) and orthologues (and syntenic) to *P. berghei* with (*). **(B)** Differential expression of AP2 TFs genes in *P. malariae* mosquito stages. Bars represent the fold change (log2) between midgut oocysts/sporozoites (red) and salivary glands sporozoites (blue). **(C)** Genomic visualization of RNA-seq coverage at differentially expressed AP2-regulated loci, H4A and histone H4B encoding genes, together with the locations of binding motifs for AP2-O4. The tracks shown are for midgut oocysts/sporozoites (red) and salivary glands sporozoites (blue) for infections 1-2. All tracks are shown at equal scale.

Four *P. malariae* AP2-TFs appear developmentally regulated in the mosquito cycle: AP2-SP2 (PmUG01_03014100, without orthologs), AP2-HC (PmUG01_12060900, ortholog to *Pb*AP2-HC, PBANKA_1319700), AP2-HS (PmUG01_12022000, without orthologs), and PmUG01_05037900 (without orthologs) (Fig 3B; Table S5). The transcriptomics profile for ApiAP2 genes in *P. malariae* mosquito stages contrasts with the pattern previously reported by *Woodford et al*. in blood stages (of mixed composition, but correlation peaking against gametocytes) (72). By reanalyzing that data, we found that the AP2 TFs AP2-P (PmUG01_09017400) and PmUG01_09024900 were the most expressed and displayed comparable peaks of expression, while gene expression levels of the AP2-TFs reported in the mosquito stages were very low in blood stages except for these two previously mentioned (Fig S3C Table S5). Looking at *P. malariae* single-cell data of mix blood stages available in the first Malaria Cell Atlas (105) (13 cells, mostly late rings and trophozoites), the TFs more expressed are AP2-O2 (PmUG01_10027900), AP2-LT (PmUG01_01029800), or PmUG01_14073400, with many others being expressed at lower levels (Fig S3D). Overall, these developmental differences between the transcriptomes support the stage-specificity of the *P. malariae* ApiAP2 family and point to a few mosquito-specific AP2-TFs playing a more prominent role during sporogony.

To study the regulatory networks of *P. malariae* AP2 TFs during infection in the mosquito, including putative binding sites and target genes, we conducted a DNA-binding motif enrichment analysis on the upstream regions (i.e., potential promoter / 5’UTR regions) for the set of genes that we categorized as highly expressed (1,005 genes, see Methods) (Table S5). For this analysis, we used as a reference *P. falciparum* AP2 TF binding sites from previous *in silico* analyses (117), but we focused on the AP2 genes that are orthologous/syntenic between *P. malariae* and *P. falciparum*. From this analysis, we obtained 3 motifs matching *Pf*AP2-O4 (TTGTTCCTATTT, TTTAGTCG, and TCTCACCT, average score ∼0.60), a motif matching *Pf*AP2-EXP (TTCCTTGTTT, score ∼0.50), and a third motif matching the uncharacterized PF3D7_1305200 (GCACGCAC, score ∼0.70) (Table S10). Out of the *P. malariae* ortholog genes, *Pm*AP2-O4 and *Pm*AP2-EXP were highly expressed in both mosquito stages, while expression for the AP2 PmUG01_14021900 (ortholog to PF3D7_1305200) was not detected (Fig 3A,Table S5).

Out of the 1,005 target genes inferred for AP2-O4 that are highly expressed, we found motif instances upstream 898 genes (Table S10B). A GO enrichment analyses for these pointed to biological processes such as movement, gliding or exit from host, sphingolipid metabolism, apoptosis, or maintenance of location. (Table S10C). Looking at developmentally regulated genes, we found motifs upstream 100 out of the 175 DEGs more expressed in oocysts/sporozoites (∼70%), and upstream 96 out of the 118 DEGs more expressed in sporozoites (∼80%). Examples of genes in this potential regulatory network include the histones H2A, H2B and H4 (DEGs more expressed in oocysts/sporozoites) (Fig 3C; Table S10B), AMA1 (DEGs more expressed in sporozoites) (Fig S4B; Table S10B), or CSP, which is highly expressed in the two stages (Fig S4A; Table S10B). Regarding AP2-EXP, in *P. falciparum* a secondary motif ([AG]C[AG]TGC[AGT]) was reported upstream > 100 sporozoite-specific genes (76). The motif TGCATGCA is also known to be recognized by AP2-EXP in multiple species (50). Looking at these two extra motifs, together with the TTCCTTGTTT reported above, we found instances upstream of 609 genes (Table S10D-E). More than half of these (332 genes) were more expressed in sporozoites (80 are DEGs), including proteins such as SPECT1, SPELD (Fig 2A; Table S10D-E), TRAP, as well as proteins involved in invasion and virulence in the liver and subsequent blood stages, such as LSAP1, P52 or UIS3 (76). When comparing the sets of potential targets to both AP2-O4 (898 genes) and AP2-EXP (609), we report overlapping, i.e. co-occurrence, for 563 genes, pointing to some degree of co-regulation. Examples of genes targeted by both include TrxL1 and ELC, which are highly expressed in both stages (Fig S4C and S5A; Table S10), whereas 335 and 46 genes appeared uniquely regulated by AP2-O4 and AP2-EXP (Fig S5B). Examples of uniquely regulated genes include PmUG01_09036800 (Fig S6A) and PmUG01_11037200 (Fig S6B), members of the lipid transfer proteins family Sec14/CRAL-TRIO and that in this study are differentially expressed genes, one more expressed in oocysts and the other in sporozoites, with predicted instances for AP2-EXP and AP2-O4 motifs, respectively (Table S10). This family has been recently characterized as crucial for lipid metabolism and host cell membrane remodeling, being potential targets for novel antimalarials (135,136).

### *P. malariae* transcriptionally variant gene families expressed in the mosquito

*Plasmodium* species possess a diverse number of gene families that appear highly polymorphic and transcriptionally variable, named clonally variant gene families (CVGs), in which a single gene can display alternative transcriptional states independently of the genotype and the stage of development. Some of these families are also multigene and display mutually exclusive expression. CVG families are key to parasite survival and are involved in a variety of processes including red blood cells invasion, metabolism, transmission, and transcriptional regulation (59,137–141). Gene expression patterns and dynamics of multigene and CVG families remain mostly unknown for *P. malariae* (14).

We first examined gene expression patterns of known clonally variant and multigene gene families as described by others (44,82), in *P. malariae* mosquito stages. Out of 1,222 genes belonging to 14 families, 123 genes appear expressed in mosquito stages in this study, 10% of the total. These included fam-m (13/283 genes), fam-l (20/396 genes), pir (11/257), stp1 (16/159), 6-cys (10/13), acs (3/4), clag (2/4), etramp (6/7), fikk (17/20), lpl (1/3), phist (12/29), rbp/rh (2/25), trag (3/42) (Table S11). Additionally, 7 out of the 123 expressed genes are DEGs, being in all cases more expressed in midgut oocysts/sporozoites, of which 5 belong to the 6-cys family and 2 to the FIKK family (Table S11). Of these 123 genes, 57 belong to the category of highly expressed in at least one infection (see Methods, 27 for midgut oocyst/sporozoites and 46 in salivary gland sporozoites) and 17 genes to the category of medium expression (all of them in salivary gland sporozoites) (Table S11). Comparing the list of known CVGs expressed in mosquitoes (123 genes) with the *P. malariae* blood-stage transcriptome reanalyzed in this study (1,463), we obtained a total of 58 matches (72) (Table S11). Of these, 5 genes are also differentially expressed in this study: 2 FIKK-encoding genes and 3 genes of the 6-cys family (Fig 4A; Table S11). It is remarkable that nine *P. malariae* CVGs, including several belonging to the aforementioned 6-cys family, contain AP2-O4 and/or AP2-EXP-associated motifs, being part of the gene regulatory network of these two transcription factors during sporogony.

**FIG 4.**
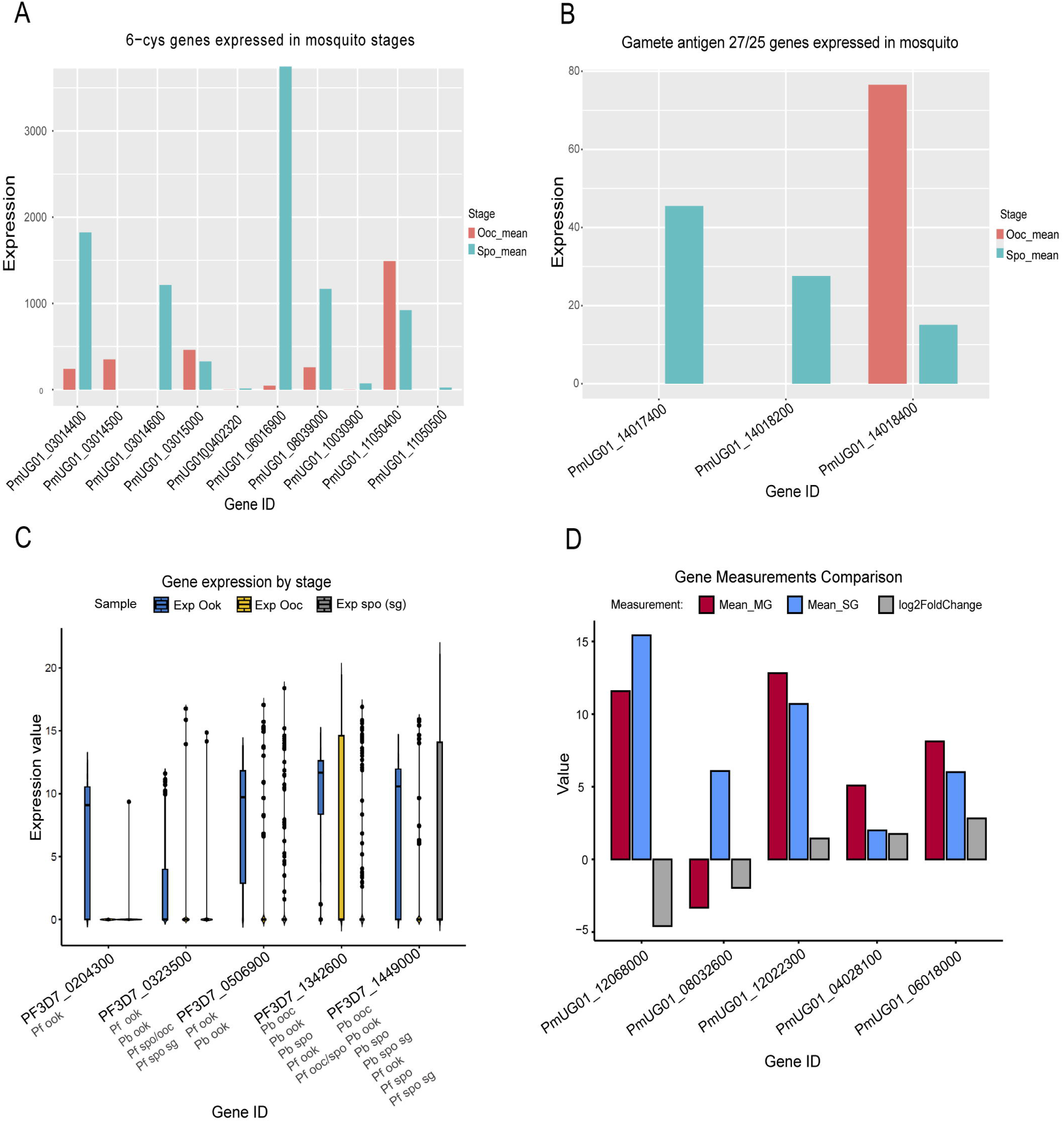
*P. malariae* clonally variant multigene families expressed in mosquito stages **(A**) Barplots showing gene expression levels for ten members of the multigene family *6-cys* expressed in mosquito stages. **(B)** Barplots showing gene expression levels for the three members of the family *gamete antigen 27/25* described as potential novel multigene family. **(C)** Barplots showing the variable and wide range of expression for 5 random HVGs computed based on *P. falciparum* scRNA-seq data. Each value corresponds to the gene expression reported in the Malaria Cell Atlas for each cell originally labeled as ookinete, sporozoites (oocysts) and sporozoites (salivary glands). **(D)** Boxplots showing for 5 random HVGs computed based on *P. falciparum* scRNA-seq data the gene expression levels in *P. malariae* midgut oocyst/sporozoites and salivary gland sporozoites samples, and the difference (log2FC).

Secondly, we aimed to identify potentially new and/or unexplored multigene families in *P. malariae* using a homology-based strategy. For this, we first searched for genes that: 1) encode apparently the same product that can or cannot be located in tandem (i.e., close to each other) and 2) show a sequence homology between gene copies that is analogous to that of the known families (∼80%, see Methods). Our homology-based strategy includes 1,396 *P. malariae* genes deemed to be potentially part of multigene families based on this search. Of these, 324 genes displayed at least one blast hit with identity >80% with any other member of its alleged family, and the vast majority of them were “Plasmodium_exported_protein”, a big family of 522 members. Out of the high confidence list of multigene candidates mentioned before (324 genes), 32 genes are expressed in *P. malariae* mosquito stages, 10 of which are highly expressed (2 in midgut oocyst/sporozoites and 8 in salivary gland sporozoites), and only 11 match ortholog and syntenic genes in other *Plasmodium* species, pointing to species-specific functions (Table S11). We also examined which of these 32 candidate genes expressed in the mosquito were also expressed in blood and obtained 13 matche*s*. Despite clear differences in expression between stages, none show statistical significance in our differential expression analyses.

The list of multigene candidates expressed in *P. malariae* mosquito stages includes members of the families MSP7-like proteins, gamete antigens 27/25, and serine repeat antigens (Table S11) More specifically, we detected that most genes coding for the gamete antigen 27/25 proteins display a higher expression in midgut oocysts/sporozoites, except for the PmUG01_14018400 gene, whose expression is significantly higher in salivary gland sporozoites (Fig 4B). As for the serine-repeat antigen genes, we identified different stage-specificity; some genes were predominantly expressed in midgut stages (PmUG01_04024400, PmUG01_04024800, PmUG01_04025200) and others with higher expression in the sporozoite stage (PmUG01_04024600, PmUG01_04024700). Particularly interesting is the case of genes encoding different variants of the MSP7-like protein, like the gene PmUG01_12030100 whose expression is variable between different infections. These results suggest a possible functional role of these candidate multigenes in different phases of parasite development within the vector.

Thirdly, to further expand the list of candidate CVGs in *P. malariae*, i.e., genes showing transcriptional variation in other species and that appear expressed in *P. malariae* mosquito stages, we analysed mosquito stages single-cell data available at the Malaria Cell Atlas for *P. falciparum* (Fig 4C) *and P. berghei* (104,105). Specifically, we identified genes showing patterns of cell-to-cell transcriptional heterogeneity that have orthologs/syntenics in *P. malariae*. Following the nomenclature by the Malaria Cell Atlas, these genes were herein considered hypervariable genes (HVGs) candidates. We applied a high-confidence approach in which only genes showing a large diverse range of expression and being identified by an approach based on standardized variance (see Methods), were considered for subsequent analysis. We analyzed independently single-cell data for *P. falciparum* developmental stages (including ookinetes, midgut oocysts and salivary gland sporozoites, see Methods). This way, a total of 484 putative HVGs candidates were identified that have an ortholog in *P. malariae* and are expressed in mosquito stages in this study (Fig 4D; Table S11). Of these, 60 genes are developmentally regulated in the mosquito (31 DEGs more expressed in midgut oocysts/sporozoites and 29 DEGs in salivary gland sporozoites). We confirmed that the majority of the 484 HVGs are highly expressed genes (being 62 highly expressed in oocyst/sporozoites, 69 highly expressed in sporozoites and 196 in both stages). These include the malaria vaccine antigen CelTOS, the Puf-family RNA-binding protein Puf2, and the pore-forming protein EXP2, merozoite TRAP family proteins: PmUG01_12028900, PmUG01_02019600 and the sporozoite traversal (GEST) protein.

From single-cell data of independent *P. berghei* developmental stages (i.e. ookinetes, oocysts, and sporozoites stages) we captured 1,896 HVGs candidates with ortholog and syntenic genes in *P. malariae* that are expressed in mosquito stages in this study. Of those, 148 genes appear developmentally regulated (81 DEGs more expressed in midgut oocysts/sporozoites and 67 DEGs more expressed in salivary gland sporozoites). Out of the 1,896 HVGs, 1,013 genes are highly expressed in our data. Several relevant examples include inner membrane complex proteins (IMC1c, IMC1e, IMC1k, ALV7, ISC1), HSP20, PSOP1, PIMMS22, gliodeosoma (GAPM1, GAPM2 GAPM3), SPATR, TRAP, GEST, P38, SPELD, CelTOS, profilin, RON2, IMP2, PPM5, PPM8, CLAMP, P36, B9, HECT1, histone(H2A), AMA1, GAMA, SSP3. Finally, a total number of 91 and 879 HVGs candidates (from *P. falciparum* and *P. berghei*, respectively) appear expressed both in blood and mosquito stages.

Altogether, the analysis of multigene and hypervariable gene families described in other *Plasmodium* spp., or proposed as such in *P. malariae* based on the comparative analysis above, resulted in a list of hundreds of putative transcriptionally variant genes that are medium or highly expressed in *P. malariae* mosquito stages. A fraction of them is also developmentally regulated. Whether they actually display clonally variant expression during *P. malariae* sporogony and which are their roles in virulence and pathogenesis, remain to be investigated in future studies.

## DISCUSSION

Many aspects of the biology of the human malaria parasite *P. malariae* remain mostly unknown (14). This study represents the first genomic study of this neglected pathogen during sporogony in the mosquito with the objective of understanding the uniqueness of its transcriptome in terms of species-specific gene repertoires, regulatory factors and virulence families having a role during this critical part of malaria parasites life-cycle.

Given the difficulty of *P. malariae* adaptation to in vitro culture, the mosquito represents an ideal experimental model that reproduces the natural conditions of an infection allowing to study parasite development and transmission and to observe biological processes that cannot be replicated in culture. Although there have been studies that approached that question by using sequential blood collections from *P. malariae* naturally infected patients, it is necessary to ensure that patients are protected from further infectious bites (142). In this context, the use of the mosquito represents a more ethical alternative to humans to study *P. malariae* infections.

To study parasite development during sporogony, is important to set the duration of the EIP in the mosquito, but in *P. malariae* this is uncertain. We chose an intermediate time point of 16 DPI to capture the parasite in sufficient numbers in midguts and salivary glands. However, this entails the risk that these samples contain a mixed population of mature oocysts and immature sporozoites. To find out, we applied a digital deconvolution approach mapping our bulk *P. malariae* RNA-seq data against recently published *P. falciparum* single-cell data of mosquito-stages and found that the highest percentage of reads correspond to the stage of development expected for each tissue, that is, oocysts in midguts and sporozoites in salivary glands. Nevertheless, the percentage of immature sporozoite reads in our midgut samples is not negligible. So, we considered this limitation in all subsequent analyses.

Another aspect that warrants caution is that in endemic areas, malaria infected Individuals often carry multiple species and distinct genotypes of the same species. In this study, we used *P. malariae* infected blood to infect mosquitoes. With regards to the presence of multiple strains/genotypes of *P. malariae*, all the infections appear monoclonal. Our initial data quality analysis reveals, however, that our samples contain, in a very high proportion, avian malaria *P. relictum* reads, whereas the remaining plasmodial reads mainly correspond to *P. malariae*. Previous studies report that, although the main vector of *P. relictum* are mosquitoes of the genus *Culex*, this parasite could eventually infect *Anopheles* mosquitoes (143,144). However, in our case, such scenario is very unlikely since the mosquitoes used in this study corresponded to laboratory reared *A. gambiae*. The only possibility for such a cross-infection is that *P. relictum* would have been already present in the human blood used in the membrane feeding assays. To date, *P. relictum* has not been documented to infect humans. A most plausible explanation is reads misassignment because of the incompleteness of the *P. relictum* reference genome, and the similarities between some species within genus *Plasmodium* (145).

In our study, 3,699 genes were identified as expressed during sporogony. This means that we successfully captured the expression of half of the *P. malariae* annotated genes (6,715). This is remarkable because of the low parasitemia characteristic of this human malaria parasite compared to other species, and the low quantity of parasite material that it is expected for mosquito stages, which could have compromised our -omics approach in terms of library quality and read coverage. Despite so, our transcriptome outperforms considerably the results of a previous study on *P. malariae* blood-stages that only detected expression of 1,463 gametocyte genes (72).

The majority of expressed genes (∼64%) with orthologs in other *Plasmodium* species conform to the expected biological functions for both oocysts and sporozoites. These include biosynthesis and metabolism in oocysts, or locomotion and symbiotic processes in sporozoites. Although only a small proportion, 7% (263), appears to be developmentally regulated based in our differential expression analysis, ∼48% (1781) of the expressed genes showed more than two-fold difference between stages but did not reach statistical significance. This result is likely due to the inherent variability between infections (different blood / parasite genotypes), as well as the limitation of the presence of mixed oocyst and sporozoite stages in the midgut noted previously.

The stage-specific expression pattern of *P. malariae* during sporogony conforms to the “just-in-time” model described for other malaria species (62), in which each developmental stage is characterized by a unique set of genes that display peaks of expression at the right moment, that is, just before the parasite needs the protein (76,132,146). An example of DEGs crucial to *P. malariae* sporogony include PIMMS22, whose ortholog in *P. falciparum* and *P. berghei* is essential for oocyst development (147). In *P. malariae* sporozoites we detected as highly expressed AMA1 which is a critical sporozoite protein in other human *Plasmodium* spp. involved in host cell invasion and motility and a key target for malaria vaccine development (148).

The expression of species-specific gene repertoires that we identified in this study is likely responsible for the unique phenotypic characteristics of *P. malariae* during infection in the mosquito. That is, approximately 20% of the *P. malariae* mosquito stages transcriptome (1,338 genes out of 6,715) is not ortholog to *P. falciparum, P. vivax* or *P. berghei*. This includes 13 of 29 ApiAP2 TFs and around 42% (78) of the developmental regulated genes. Some examples are ASP, TLP, EXP2, CLAMP and TRAP which are known to be involved in processes related to species-specific differences. For example EXP1 and EXP2, involved in different red blood cells preference, cytoadherence or persistence, between *P. vivax* and *P. falciparum* (46,149) .

Despite of the advances in the knowledge of the genome of this organism (44), 20% of all genes (1,320) still do not have an annotated function. In this study we report expression of 70% of the non-annotated *P. malariae* genome (955). Importantly, around a third of the DEGs are genes with unknown functions (82 out of 263). We attempted to annotate expressed genes lacking an associated gene description, using a tool previously tested in *Trypanosoma brucei* for the same purpose (115), resulting in 136 genes with newly annotated GOterms. Further experimental studies are now needed for complete functional characterization.

In *Plasmodium*, the family of ApiAP2 TFs are essential master regulators which various members playing a role at distinct stages of the parasite development (49,117,121). Knowledge of this family in *P. malariae* is limited to a single study in blood stages (72). They report the expression of only two AP2 members, *Pm*AP2-P and PmUG01_09024900. In this study, we described the expression patterns of the entire family (29 members) during *P. malariae* sporogony. Most of the PmAP2 TFs (23) appear to be expressed in the mosquito, 14 of them appeared more expressed in midgut oocyst/sporozoites and 9 in salivary gland sporozoites, however only 4 *Pm*AP2 were significantly differentially expressed.

Out of the 23 PmAP2 proteins expressed in mosquito stages, 3 and 9 members, respectively, are ortholog and syntenic to *P. falciparum* and *P. berghei* AP2s. Two examples are AP2-O4 and PmUG01_09024900. Previous studies have reported that PbAP2-O4 is only expressed in sexual stages, playing a more prominent role in the oocyst stage (50,121). With regards to PmUG01_09024900, early work reported the expression of its ortholog PF3D7_1115500 in *P. falciparum* mosquito stages (75,76). Also highly expressed in *P. malariae* mosquito stages and ortholog and syntenic to *P. berghei* AP2s, are AP2-I, PmUG01_09017400, AP2-HC (also developmentally regulated), AP2-G3 (previously AP2-FG). PfAP2-I is a key regulator of red blood cell invasion in *P. falciparum* (150), but is not ortholog to *Pm*AP2-I, suggesting an unique a new role of this TF in the mosquito cycle. Noticeably, it has been described that AP2-G3 regulates gene networks involved in *P. berghei* female gametocyte maturation, fertilization and zygote development (151). Finally, *Pm*AP2-EXP is ortholog (and syntenic) to *Pf*AP2-EXP, and has been proposed to be a master regulator of *P. falciparum* sporogony (76). Altogether, for AP2 proteins with orthologs/syntenics in other Plasmodium spp., our findings reveal several similarities across species but also point to new functionalities of these proteins in *P. malariae*.

One remarkable finding of this study is the fact that most of the *P. malariae* AP2 TFs that we identified as highly expressed in mosquito stages, don’t have any ortholog in *P. falciparum* or in *P. berghei.* This is the case of AP2-O, AP2-TEL, AP2-L with peaks of expression in midgut oocyst/sporozoites and PmUG01_05037900, AP2-L, AP2-O and AP2-O5 overexpressed in salivary glands sporozoites. The same is true for 3 out of 4 developmentally regulated PmAP2: PmUG01_05037900, AP2-SP2 and AP2-HS.

Apart from the differences between species, the transcriptome of *P. malariae* during sporogony contrasts markedly with the pattern reported for blood stages, where only AP2-P and PmUG01_09024900 appeared expressed. These two are also highly expressed in mosquito stages but together with 21 other AP2 proteins, 11 of them being *P. malariae* specific. The reduced number of AP2 transcription factors reported for the blood cycle is consistent with the lower number of expressed genes observed in that study, which may be related to the coverage and quality of the available transcriptome. In either case, these results highlight the differences in the regulation and function of the *P. malariae* ApiAP2 family of TFs across the parasite life-cycle.

To learn more about the gene regulatory networks of ApiAP2 transcription factors in *P. malariae*, we focus on those members of the family that appear expressed and have orthologs and syntenics in *P. falciparum*. Only AP2-O4 and AP2-EXP meet this requirement. We found enriched motifs matching the binding sites for AP2-O4 in 898 highly expressed genes being 431 in midgut oocysts/sporozoites and 467 in salivary gland sporozoites and at 196 out of the 263 differentially expressed genes. Our results agree with the key role of AP2-O4 in rodent malaria parasites being essential for the oocyst formation (50,121). Similarly, considering the three existing motifs for PmAP2-EXP we found instances at the promoters of 609 highly expressed genes, 332 in oocyst/sporozoites and 277 in sporozoites. AP2-EXP had already been implicated in transcriptional regulation during sporogony both in *P. falciparum* and *P. berghei* (49,76,80,121,152). In agreement, we identified target genes like SPECT1 and TRAP, both crucial for parasite motility, tissue invasion in the sporozoite stage and, consequently, in the transmission of malaria in other species of *Plasmodium* (153,154). We also report target genes related to the subsequent invasion of hepatocytes. Examples of these genes are P52, responsible for the formation of a parasitophorous vacuole required for parasite replication in the hepatic stage (73), or UIS3 that is also essential for parasite survival in the hepatocytes (155). These findings suggest that these AP2 TFs could play similar functions during *P. malariae* development in the mosquito and in humans. Also, it is important to note the number of instances that both TFs have in common representing ∼60% of the instances reported. This would point to a certain degree of co-regulation. So far, there are no studies indicating that these two TFs work together even in their *P. falciparum* orthologs.

Collectively, the pattern of stage-specific expression that we report for P. malariae ApiAP2 TFs, the gene regulatory networks inferred, and the similarities/differences with other species, support the role of these proteins as key regulators of *P. malariae* development and life-cycle progression in the mosquito. Chromatin studies, like ChIP-seq and ATAC-seq, accompanied by protein function validation, would be now needed to further characterize these *P. malariae* master regulatory proteins.

Clonally variant gene families are essential for parasite survival via processes such as antigenic variation or immune evasion, red blood cells invasion and sexual differentiation (82). Mutually exclusive expression is often associated with multicopy gene families such as the *var* gene family in *P. falciparum*, or the *pir* family, which is the most ubiquitous multigene family together with *stp1* in *P. malariae*. In this species, there are no orthologs to the *var* gene family, the *pir* gene family is very restricted (∼50% are pseudogenes, compared to 9% in *P. vivax* or 25% in *P. o. curtisi*), and *stp1* displays a species-specific expansion, together with the gamete antigen 27/25 or merozoite surface proteins (*msp3*, *msp7*) (44,54,82,142). Contrary, several *P. malariae*-specific multigene families have been newly described in the latest genome assemblies, including some that appear to be expanded when compared to other species. This is the case of two novel large multigene families, *fam-l* and *fam-m*, that do not have orthologs in other species, but their structure overlaps 100% with the *P. falciparum* protein RH5, a RNA binding protein essential for erythrocyte invasion (44,54). Importantly, the transcriptional profiles of these multigene families in *P. malariae* remain poorly known, partly due to the paucity of genomic and transcriptomic data for this species, but also because it is very challenging to analyze their expression because most genes tend to be expressed at low levels, are located in the sub telomeres and in highly repetitive regions.

Remarkably in this study we detected expression of 123 members of these CVG families, 57 of them being highly expressed, including: *fam-l*, *fam-m*, *pir*, *stp1*, tryptophan-rich antigens, 6-cys proteins and early transcribed membrane protein, among others. Only 7 genes appear differentially expressed between the two developmental stages, most of them belonging to the 6-cys family, P36, P52, PmUG01_03014600, P38 and B9. One example is the 6-cys family, in which proteins like P52, P36, P38, P12p, P41 and PSOP12 are overexpressed in *P. malariae* sporozoites. The literature describes that the complex formed by P36 and P52 has a role in *P. yoelii* sporozoites, during establishment of the parasitophorous vacuole and in the invasion of hepatocyte (156). Similarly, P38, P41 and P12p have roles during invasion of RBC in *P. vivax* and *P. falciparum* (157–159), while PSOP12 in is reported to be expressed in *P. falciparum* gametocytes, ookinetes and oocyst and in *P. berghei* affects oocyst production (157). Whether these proteins play similar roles during *P. malariae* sporogony, it is not known.

Interestingly, 9 CVGs *P. malariae* genes, including several members of the 6-cys family discussed above, have motif instances for AP2-O4 and/or AP2-EXP suggesting they are target genes of these two TFs during sporogony.

Contrary to our results in mosquito stages, a previous RNA-seq study in *P. malariae* gametocytes did not find expression for most multigene families, including *fam-m*, *fam-l* and *pir* (72). That study suggested that this pattern is because these families do not have a function during the IDC, as opposed to other exported proteins, such as *stp1* or *phist* (72).

Apart from these well-known families, one assumption of our study is that there might be other potentially new and/or unexplored multicopy gene families in *P. malariae*. Indeed, we provide a list of 324 candidates based on sequence similarity, and of these we report the expression of 32 genes, 10 of them being highly expressed. Apart from being expressed in mosquito stages, our reanalysis of published gametocytes/asexuals *P. malariae* transcriptomic data revealed that these families are also expressed in the blood cycle. These genes correspond mostly to serine-repeat antigens, one MSP-7 like proteins, and gamete antigens 27/25. The *P. malariae* genome includes 20 copies of gamete antigens 27/25, compared with a single copy in other examined *Plasmodium* species (44). It has been shown that disrupting this single gene in *P. falciparum* impacts membrane integrity during gametocyte maturation (54). Members of another family here identified as potentially multicopy in *P. malariae*, the merozoite surface protein 7 (*msp7-like*), appeared highly expressed in *P. malariae* mosquito stages. These proteins have previously been suggested to be important factors associated with the *P. vivax* preference for younger red blood cells and with immune evasion (46,160) and it is proposed that the differences in transcriptional patterns are due to different functions throughout the *P. vivax* cycle (160). *P. malariae* can also produce persistent infections and it is possible that this type of antigenic variation would permit prolonged and latent *P. malariae* infections, as described for other species.

Another novelty of our work has been the use of Plasmodium spp. published single cell data to propose hypervariable genes in *P. malariae*. In doing so, we generate a list of 2,380 candidate genes which show cell-to-cell transcriptional heterogeneity in *P. falciparum* and/or *P. berghei* and have orthologs in *P. malariae.* Approximately 63% (1,497) of these HVGs candidates based on other species evidence are highly expressed in *P. malariae* mosquito stages and 60 are developmentally regulated. We also report that 20-40% (from *P. falciparum* and *P. berghei*, respectively) appear to be expressed in *P. malariae* blood stages. Several relevant examples include the malaria vaccine antigen CelTOS, TRAP or the gamete egress and sporozoite traversal (GEST) protein. CelTOS is expressed in both ookinetes and sporozoites of several *Plasmodium* species. It is essential for the parasite’s ability to traverse host cell barriers, facilitating invasion into mosquito midgut cells and human hepatocytes (132,161). *Pb*GEST is involved in the egress of gametes from red blood cells and the traversal activity of sporozoites (162). The expression of this protein in *P. berghei* and *P. falciparum* was previously reported to be more expressed in response to environmental cues within the mosquito host, underscoring its role in parasite development and transmission (132,162).

Collectively these analyses result in a list of putative novel clonally variant families in *P. malariae* showing evidence of high expression in mosquito and blood stages. Despite we report variable expression for many of these candidates in *P. malariae*, not only between stages but also across different infections, it is important to note the limitation of using natural infections to study the CVG families. That is, variable expression can be confounded by the presence of multiple genotypes and of mixed stages in the mosquito midgut. Future work is needed, for example at the level of single cell transcriptomes on developing oocysts and sporozoites, to confirm transcriptional heterogeneity and mutually exclusive expression happening in *P. malariae*.

In conclusion, we report for the first time the transcriptome of *P. malariae* during sporogony in the mosquito filling an important gap related to our current knowledge on the biology of this neglected human malaria parasite. We characterized stage-specific transcriptional programs that recapitulate the different biological functions for each developmental stage during sporogony and conform to what has been described for other malarias. However, our analyses reveal species-specific gene repertoires expressed in mosquito stages which are likely responsible for unique life-history traits of this species. We could also gather valuable data on the transcriptional dynamics and role of *P. malariae* ApiAP2 family of transcription factors including some insights into their gene regulatory networks during sporogony. Finally, we describe the expression in the mosquito of known *P. malariae* CVG families and putative novel hypervariable genes and multigene families with possible roles in parasite virulence and pathogenesis. All these insights are crucial to understanding what makes these pathogens more or less virulent and transmissible, contributing to designing more innovative strategies for malaria eradication.

## DATA AVAILABILITY

The RNA-seq data generated in this study has been deposited in the GEO database under accession number GSE231944. All processed datasets supporting the conclusions of this work are available in the supplementary materials.

## ACKNOWLEDGEMENTS

We are grateful to all the volunteers from Burkina-Faso that kindly donated blood and to all the technicians from INSTech that helped us during the blood-donors recruitment, experimental infections and dissections. We also thank Amparo Hidalgo for technical support. This manuscript is part of the doctoral thesis of Bárbara Díaz Terenti in the Ph.D. program of molecular biology and biochemistry of the University of Granada.

## LEGENDS

**Supplementary Fig S1.**

**(A)** Correlation plot showing similarity between infections (I1, I2) and developmental stages (*P. malariae* oocysts and sporozoites). Samples are ordered by an unsupervised hierarchical clustering approach and clustered by stage rather than biological replicate (experimental infection) both immediately after alignment to reference genome (top) and after processing and computing read counts (bottom). **(B)** Principal Component Analysis on the biological replicates (I1, I2) and developmental stages (midgut oocysts/sporozoites and salivary glands sporozoites). The plots show the percentage of variance explained by PC1 and PC2 axes and the distribution of the samples along the first two Principal Components. Samples clustered by stage rather than biological replicate. **(C)** Correlation plot showing the similarity between the *P. malariae* samples in this study and previously published transcriptomes during sporogony for *P. falciparum* (75,76) , *P. vivax* (73) and for *P. berghei* from *PlasmoDB* (91). Expression data are TPM-normalized gene counts. The pairs with significant correlation coefficients are indicated. The peaks of correlation between matching developmental stages in the different species are highlighted in red. **(D)** Barplots showing the deconvolution of the *P. malariae* bulk RNA-seq data samples in this study using previously published *P. berghei* single-cell data (105). Matching stages account for the larger proportions.

Supplementary Fig S2.

**(A)** Barplots showing the number of SNPs in the sequences when comparing with the reference genome. The differences between infections for the same developmental stage point to the multiclonality of independent experimental infections. **(B)** GO terms obtained in the gene ontology enrichment analysis performed on the 118 differentially expressed genes (DEGs) that appear more expressed in salivary glands sporozoites. **(C)** GO terms obtained in the gene ontology enrichment analysis performed on the 145 differentially expressed genes (DEGs) that appear more expressed in midgut oocyst/sporozoites.

Supplementary Fig S3.

**(A)** Word cloud showing GO enrichment for the 326 “no function” genes without orthologs in *P. falciparum, P. berghei* and *P. vivax*. **(B)** Word cloud showing GO enrichment for the 388 “no function” genes with orthologs in *P. falciparum, P.berghei* and *P. vivax,* but without GO terms associated. **(C)** Expression patterns of *P. malariae* AP2 TFs in blood stages, data from *Woodford et al.* (72) **(D)** Expression patterns of *P. malariae* AP2 TFs in blood stages, data from the malaria cell atlas (105).

Supplementary Fig S4.

**(A-C)** Genomic visualization of expressed AP2-regulated loci. Examples of genomic regions include differentially and highly expressed genes CSP (A), AMA1 (B), and TrxL1 (C), together with the positions of binding motifs for AP2-O4 and AP2-EXP. The tracks shown are for midgut oocysts/sporozoites (red) and salivary glands sporozoites (blue) for infections 1-2. All tracks are shown at equal scale.

Supplementary Fig S5.

**(A)** Genomic visualization of expressed AP2-regulated loci. Examples of genomic regions include differentially and high expressed gene ELC, together with the positions of binding motifs for AP2-O4 and AP2-EXP. The tracks shown are for midgut oocysts/sporozoites (red) and salivary glands sporozoites (blue) for infections 1-2. All tracks are shown at equal scale. **(B)** Venn diagram showing overlapping AP2-EXP and AP2-O4 motifs.

Supplementary Fig S6.

**(A-B )** Genomic visualization of expressed AP2-regulated loci. Examples of genomic regions include differentially and highly expressed gene PmUG01_09036800 (A) and PmUG01_11037200 (B), together with the positions of binding motifs for AP2-O4 and AP2-EXP. The tracks shown are for midgut oocysts/sporozoites (red) and salivary glands sporozoites (blue) for infections 1-2. All tracks are shown at equal scale.

**Supplementary Table S1.** Mosquito experimental infections.

Two experimental infections (I1-I2) were performed. Microscopic confirmation of midgut infection including the prevalence (percentage of infected mosquitoes), the intensity of infection (median number of oocysts), the numbers of mosquitoes and the range of oocyst numbers.

**Supplementary Table S2.** Taxonomic classification of the sequencing reads.

**(A-E)** Taxonomic classification and species composition of the reads for each sequencing lane, experimental infection and assayed mosquito tissue (midguts, salivary glands), respectively.

**Supplementary Table S3.** Read alignments statistics. Summary of RNA-seq alignment and quality control statistics.

**Supplementary Table S4**. Bulk RNA-seq deconvolution.

**(A)** Relative cell-type composition of the *P. malariae* samples in this study based on available *P. falciparum* single-cell RNA-seq profiles (125). **(B)** Relative cell-type composition of the *P. malariae* samples in this study based on available *P. Berghei* single-cell RNA-seq profiles from the Malaria Cell Atlas (105).

**Supplementary Table S5.** RNA-seq gene expression levels.

Gene expression (DESeq2-normalized RNA-seq counts) for the *P. malariae* transcriptome. Columns include ranking by expression (high, medium or low), functional annotation, the ratio of expression between experimental infections, orthology information and the gene expression levels at gametocytes based on our reanalysis of previously available data (72).

**Supplementary Table S6.** Gene ontology terms overrepresentation analysis differentially expressed genes.

Gene ontology (GO) terms overrepresentation analysis for the sets of stage-specific DEGs. Results include the biological processes (BP), molecular functions (MF) and cellular components (CC) GO terms, and the KEGG and MetaCyc metabolic pathways used by PlasmoDB (91).

**Supplementary Table S7.** Gene ontology terms overrepresentation analysis of genes without orthologs in *P. falciparum, P. berghei* and *P. vivax*. Results include the biological processes (BP), molecular functions (MF) and cellular components (CC) GO terms used by PlasmoDB (91).

**Supplementary Table S8.** List of expressed genes without a product description *“no function”*. Gene expression values are DESeq2-normalized RNA-seq counts. Columns correspond to functional annotation, orthology information and GO terms obtained by PlasmoDB (186) or by PANNZER2 (113).

**Supplementary Table S9.** Gene ontology terms overrepresentation analysis of *P. malariae “without annotated function”* expressed genes (GO terms obtained from PlasmoDB (91) (BP and CC). TopGO overrepresentation analysis in genes without orthologs in *P. falciparum, P. berghei* and *P. vivax* (GO terms predicted by PANNZER2 (113)) also performed in genes with orthologs *P. falciparum, P. berghei* and *P. vivax* (GO terms obtained by PANNZER2).

**Supplementary Table S10.** Motif occurrences in mosquito stages highly expressed genes.

**(A)** List of enriched consensus motifs corresponding to known ApiAP2 TFs in the promoters of highly expressed genes in mosquito stages **(B)** Instances corresponding to AP2-O4 motifs located at the promoters of highly expressed genes. **(C)** GO terms overrepresentation analysis of the highly expressed genes with AP2-O4 motif instances at promoters. **(D-E)** Instances corresponding to AP2-EXP motifs obtained in this study (D) and by others (E) that are located at the promoters of highly expressed genes (118). **(F)** GO terms overrepresentation analysis of the highly expressed genes with AP2-EXP motif instances at promoters.

**Supplementary Table S11.** Expression of clonally variant multigene families and hypervariable gene families.

List of expressed genes belonging to known CVGs previously described, potentially CVGs and HVGs obtained by single-cell data. Columns include expression, ranking by expression (high, medium or low), functional annotation, orthology information and production method.

