## Supplementary figures and images for "The dynamic transcriptome of *Plasmodium malariae* in the mosquito"

### Supplemental figure 1

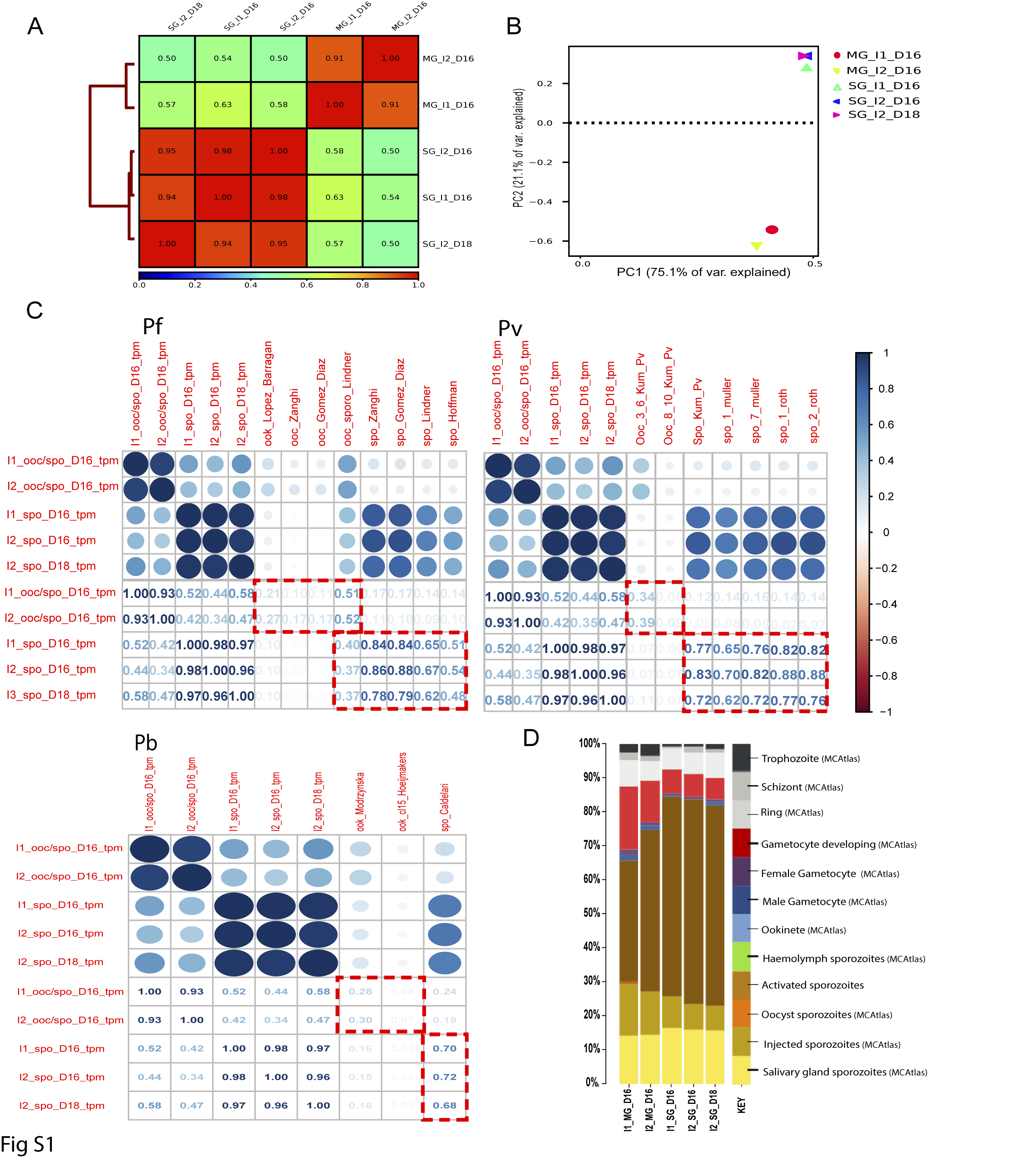

### Supplemental figure 2

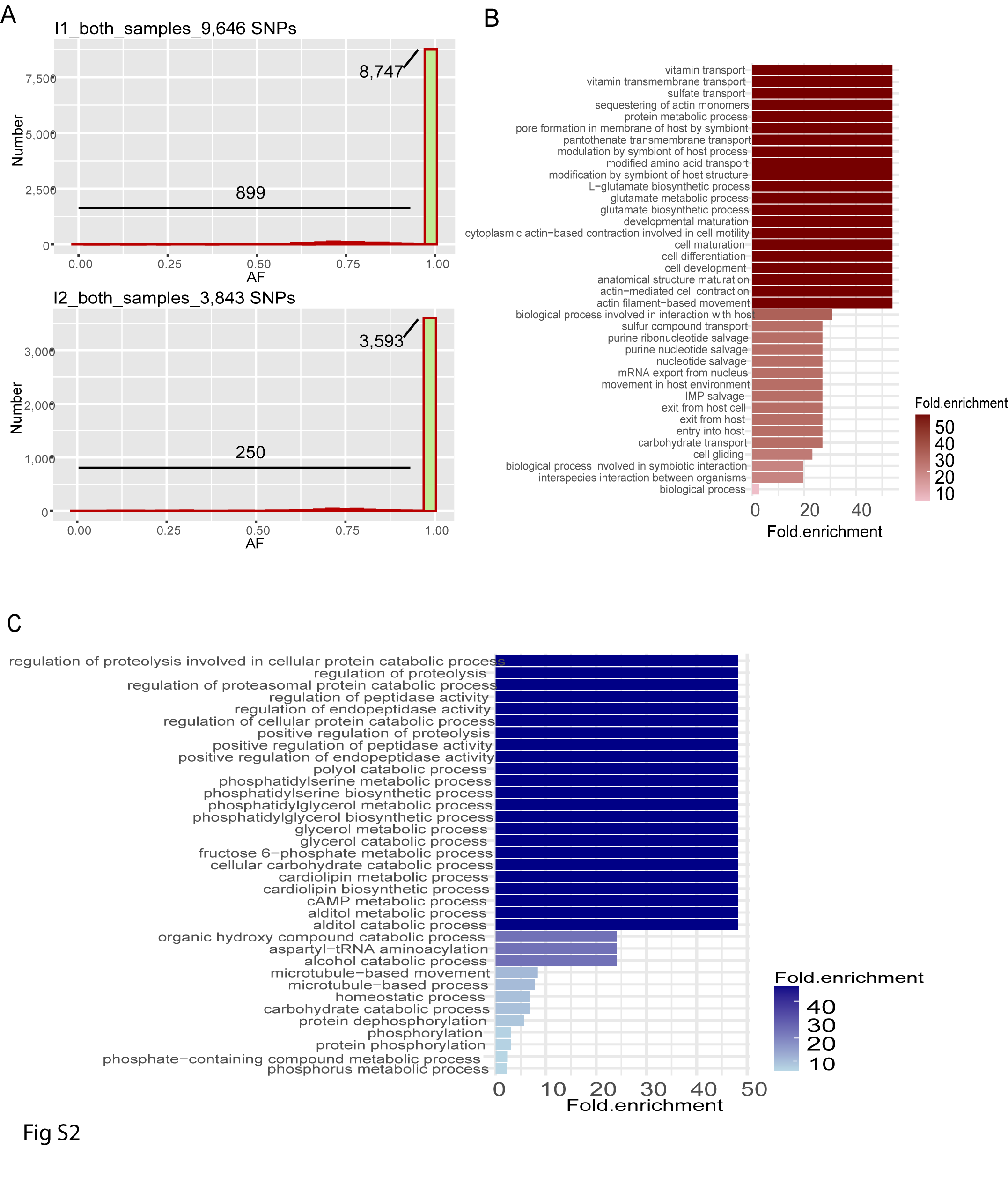

### Supplemental figure 3

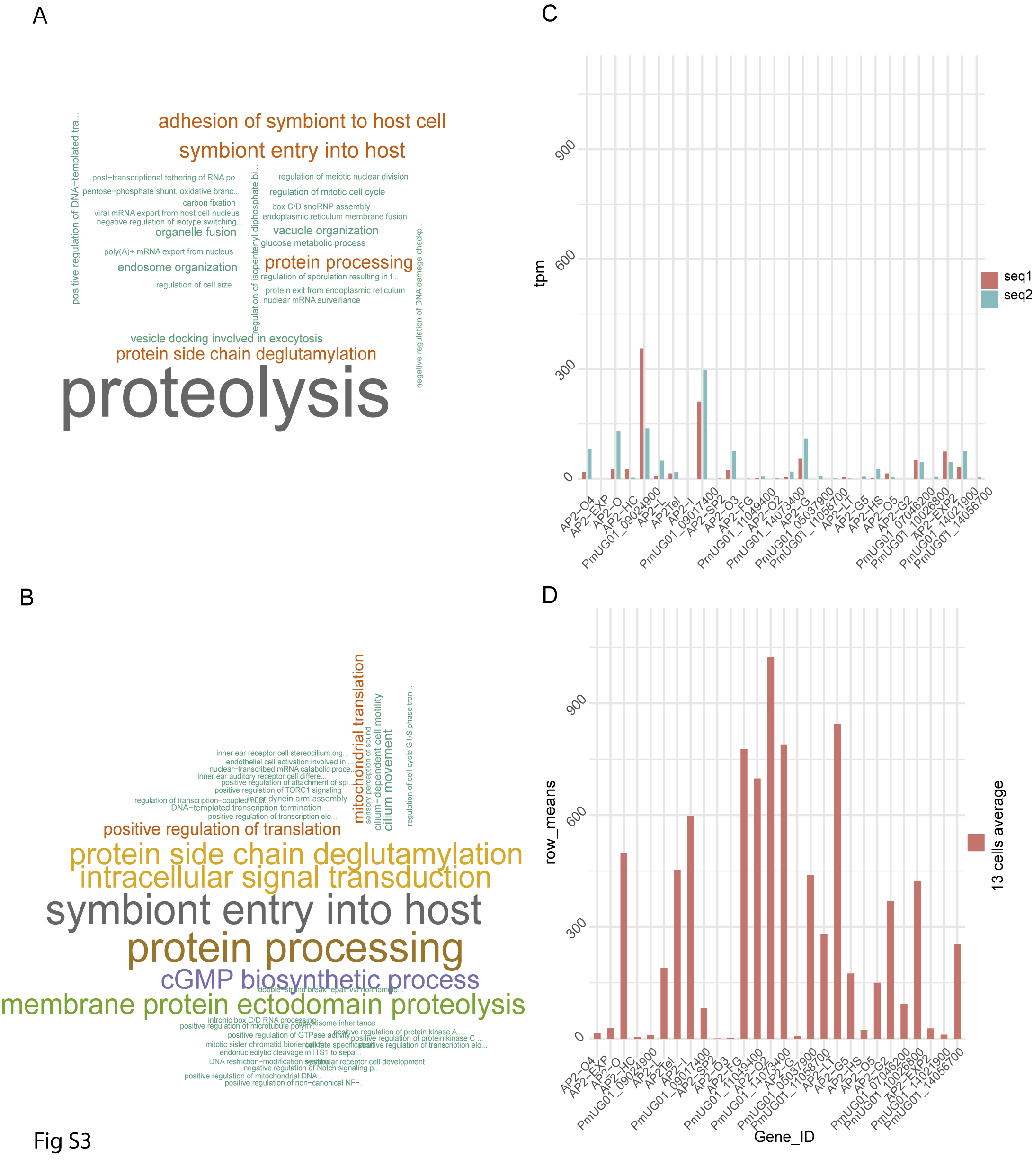

### Supplemental figure 4

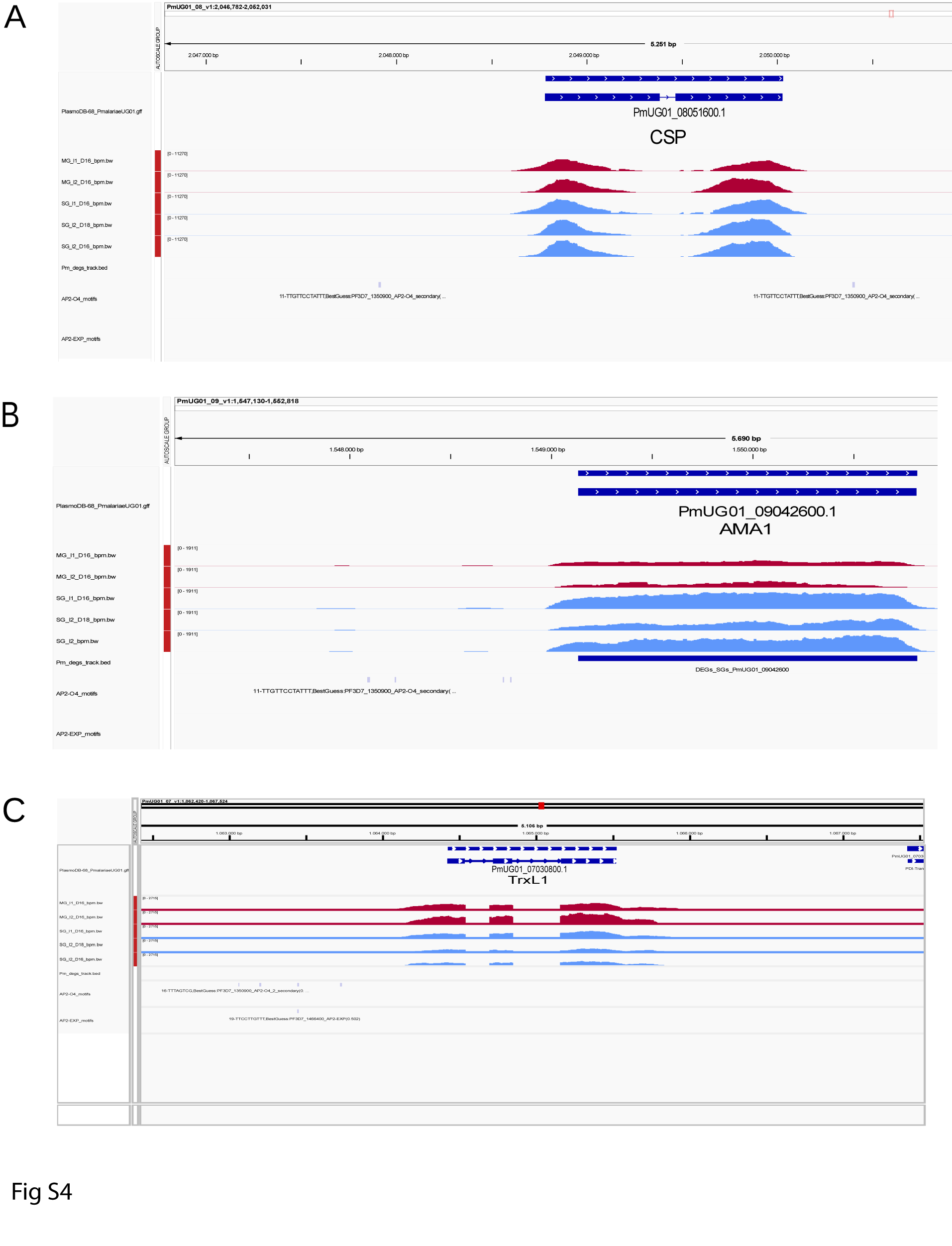

### Supplemental figure 5

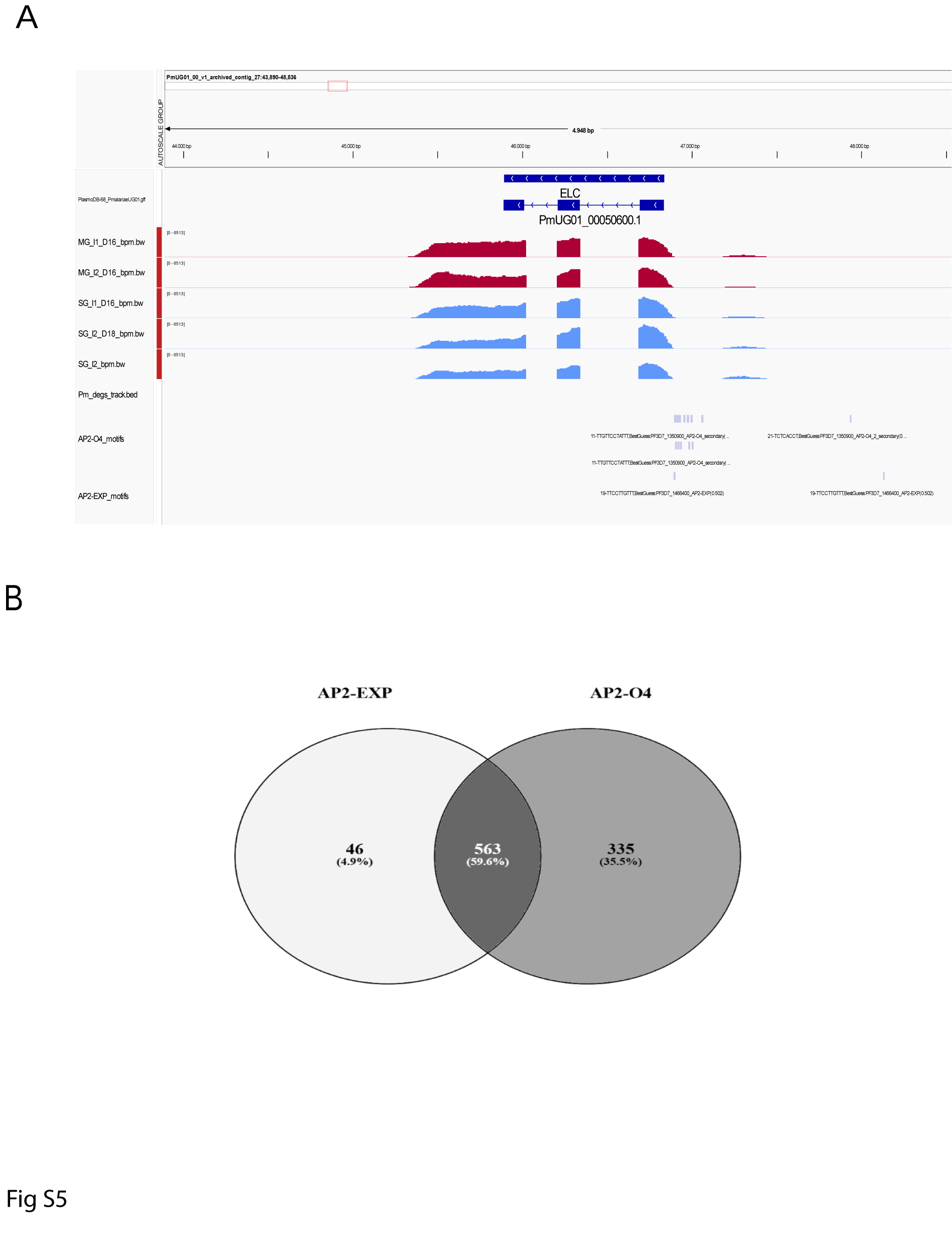

### Supplemental figure 6

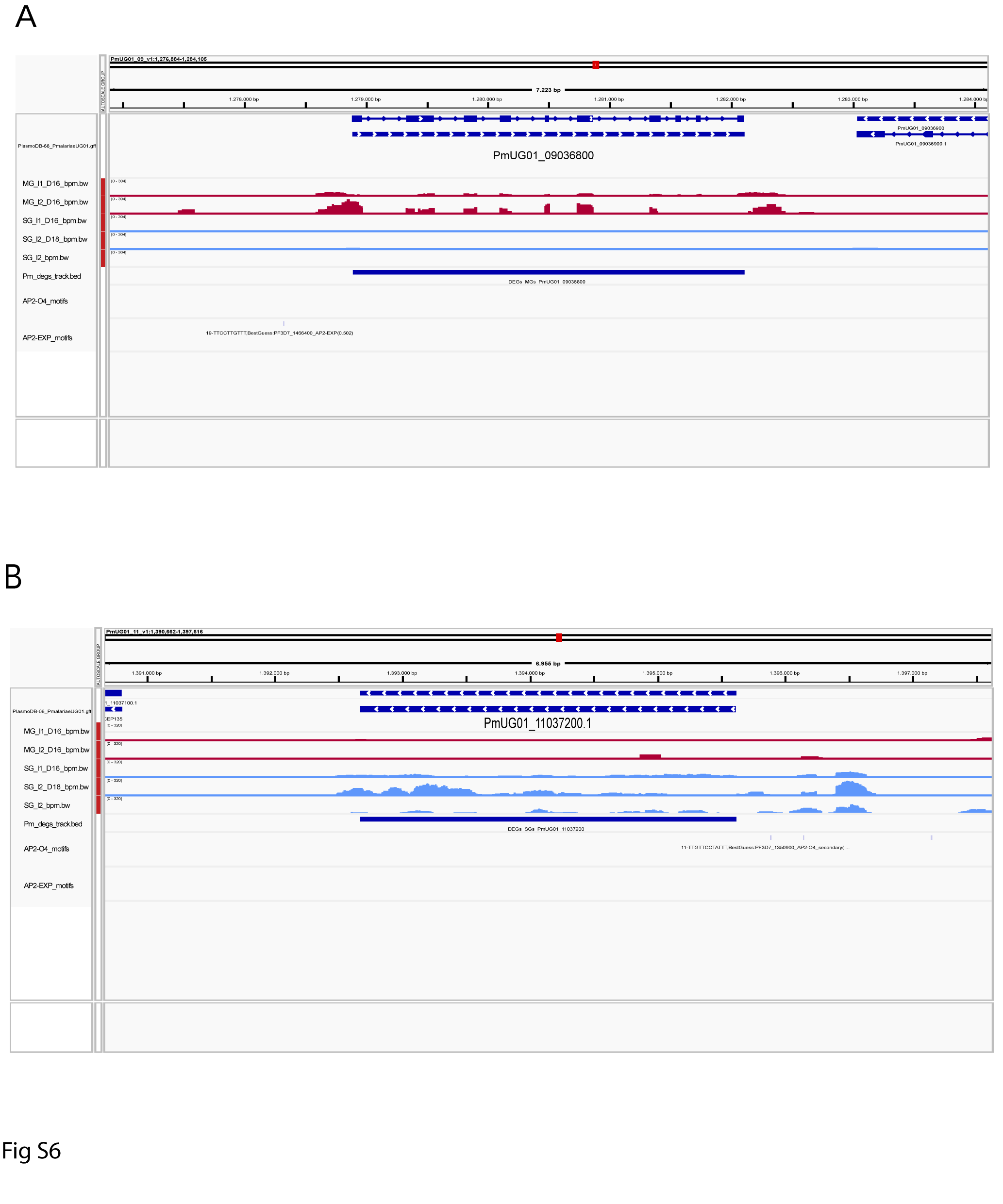
